# Coupled transcriptomic divergence establishes a human-specific synaptic glial precursor state

**DOI:** 10.64898/2026.08.19.745872

**Authors:** X D. Sheu, Yuki Y. Yamauchi, Rintaro Amano, Yosuke Nakano, Jiro Yoshino, Ikuo K. Suzuki

## Abstract

The mammalian cerebral cortex is built from a conserved developmental program, yet exhibits profound species-specific complexity. To decode the regulatory changes driving human brain evolution, we reconstructed and aligned continuous single-cell differentiation trajectories across the developing human, macaque, mouse, and ferret cortices. This comparative framework revealed a fundamental principle of transcriptomic evolution during mammalian cortical development: while stable expression is the mammalian default, genes that diverge strictly shift their allocation to cell differentiation trajectories and developmental timing in tandem. By isolating these coupled regulatory shifts to the human lineage, we revealed that a canonical synaptic gene network uniquely redeployed into early human oligodendrocyte precursor cells (OPCs). Human, chimpanzee, and gorilla cortical organoids confirmed that this neuron-like OPC state is an exclusively human innovation. Spatial transcriptome analysis found that these specialized OPCs engage adjacent neural progenitors (outer radial glia) via synaptic-adhesion signaling during neurogenetic period. These findings demonstrate that this coupled spatiotemporal rewiring establishes novel developmental microenvironments, providing a discrete molecular engine for human cortical evolution.

## Introduction

The mammalian cerebral cortex is constructed through a conserved developmental program, yet the resulting complexity differs enormously across species. The human cortex, for instance, contains approximately three orders of magnitude more neurons than the mouse cortex^1^. Nevertheless, the temporal order of developmental events and cell differentiation trajectories remains highly stereotyped and conserved: a common primary neural progenitor, the ventricular radial glia (vRG), sequentially generates deep-layer neurons, superficial-layer neurons, and finally glia^2–4^. Between the primary vRG progenitor and terminally differentiated cells, specific intermediate progenitor types regulate lineage trajectories and amplify descendant populations in a species- or phylogeny-dependent manner. For example, large-brained animals, including humans, possess an additional self-renewing intermediate progenitor population, such as outer radial glia (oRG). While the expression profile of oRGs closely resembles that of vRGs, they reside in a distinct spatial domain within the fetal cortex ^5–7^. Owing to the massive neurogenic and gliogenic capacity of oRGs, human cortical development features an extended period of late neurogenesis that overlaps with gliogenesis. Thus, humans inherit a conserved cellular differentiation program that has undergone notable spatiotemporal modifications to ultimately yield an extraordinarily elaborate neural circuit. How these conserved cellular trajectories give rise to such divergent neural architectures, and what specifically distinguishes the human lineage, remains a central question in cortical evolution.

While a few human-specific gene duplications (e.g., *NOTCH2NL*^*9,10*^, *ARHGAP11B*^*11,12*^) directly expand progenitor populations, the vast majority of the genome is shared across mammals. Comparative work has therefore converged on the view that most species differences arise from changes in how a conserved gene inventory is deployed rather than from changes in its content^13–16^. Known human or primate specific modifications include the retention of *PAX6* expression in basal progenitors in oSVZ^17,18^, the elevated expression of the Wnt receptor *FZD8* in progenitors^19,20^, and the protracted schedule of neuronal maturation^21^. In each case, a conserved gene is used in a different cell population, at a different point in development, or both.

Systematically identifying such evolutionary modifications requires a reliable reference. Because a developing and differentiating tissue is a continuum, matching species by discrete chronological stages or conventional cell-type markers often obscures continuous transcriptomic changes^22–27^. Reconstructed differentiation trajectories offer a robust alternative. By mapping how cell populations shift from one recognizable state to another, we can align these continuous routes of transcriptomic change across species to systematically compare how the shared genome is deployed.

Here, we reconstruct and align developmental trajectories from single-cell atlases of the developing human, rhesus macaque, mouse, and ferret cortex^28,29,30,31,32,33^. Applying this comparative framework genome-wide revealed a fundamental principle of transcriptomic evolution. While broad and conserved expression across differentiation lineages is the default state in mammals, orthologs that undergo evolutionary divergence consistently shift their trajectory usage (allocation) and developmental timing (progression) in tandem. By isolating these coupled regulatory shifts specifically on the human phylogenetic branch, we identify a uniquely human precursor state: an early population of oligodendrocyte precursor cells (OPCs) defined by the transient expression of neuronal synaptic genes. Located in the outer germinal zone, these specialized, neuron-like OPCs interact directly with the outer radial glia pools that characterize human neurogenesis.

## Results

### A single framework yields comparable developmental trajectories in four mammals

Cortical development proceeds through a branching hierarchy, in which ventricular radial glia (vRG) give rise to progenitor intermediates and ultimately to neurons, astrocytes, and oligodendrocytes (**Figure 1A**). To compare this process across mammals, we sought to reconstruct these differentiation routes in each species and to define them equivalently. Two forms of evolutionary change in expression of a given gene could be distinguished: a “trajectory change”, in which route an ortholog is expressed on, and a “progression change”, in when along a route it is expressed (**Figure 1B**). Both measures depend on the routes being defined in the consistent manner in every species, and we therefore began by assembling a comparable developmental substrate. Single-cell transcriptomic atlases were collected from human^28^, rhesus macaque^29^, mouse^30,31^ and ferret^32,33^. Developmental stages were matched across species using histologically defined landmarks, as described previously^34–40,41–44.^ Each atlas was then restricted to a fixed developmental window: gestation week 10 (GW10) to GW36 in human, embryonic day 37 (E37) to E110 in macaque, E10 to postnatal day 4 (P4) in mouse, and E25 to P10 in ferret (**Figure 1C**). These windows cover neurogenesis and the majority of gliogenesis in all four species.

**Figure 1.**
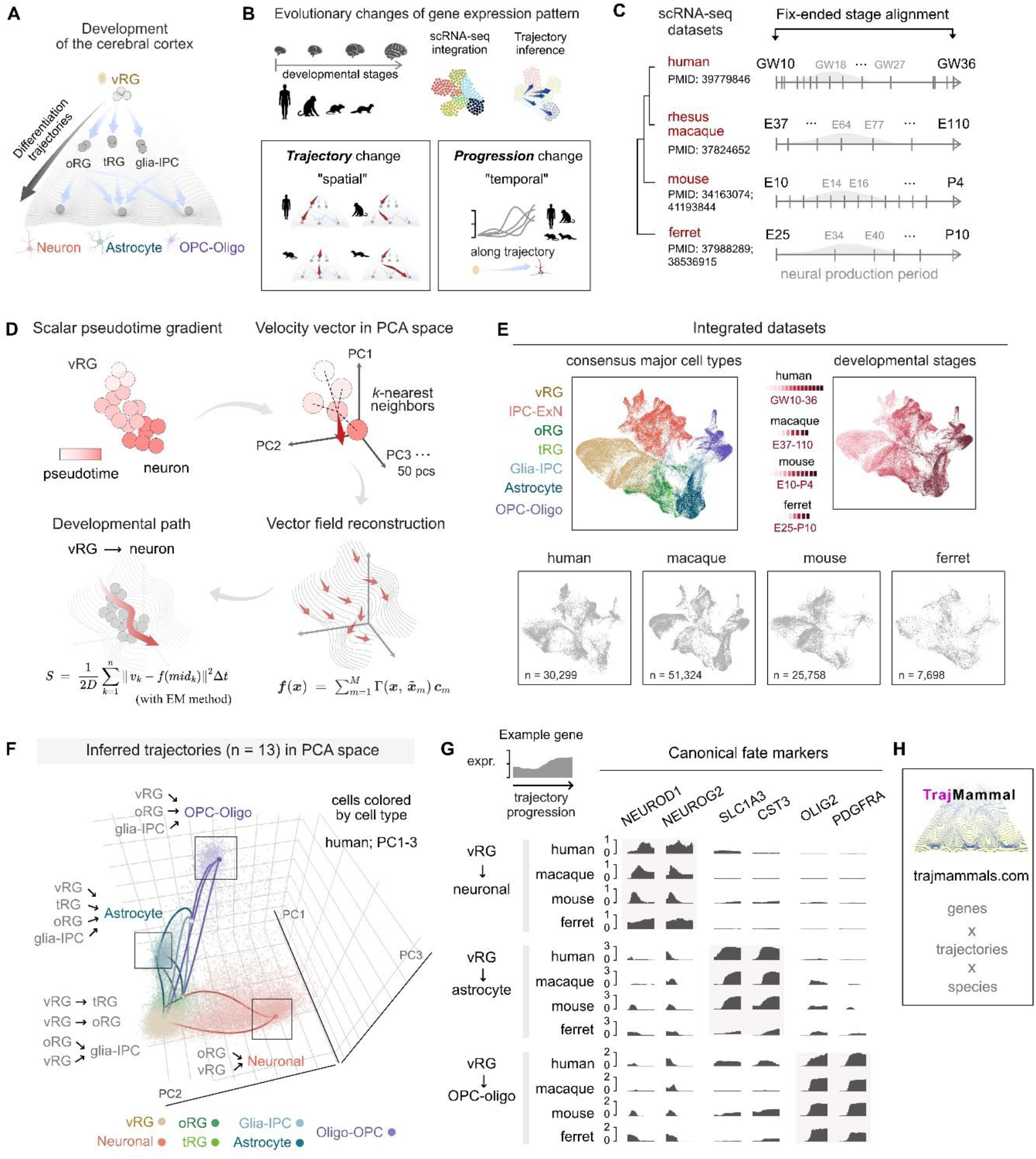
Reconstruction of single-cell developmental trajectories in four mammalian cortices. **A.** Cortical development as a branching hierarchy. Ventricular radial glia (vRG) give rise to intermediate progenitors (outer radial glia or oRG, truncated radial glia or tRG, glia intermediate progenitor cells or glia-IPC) and ultimately to neurons, astrocytes, and oligodendrocytes. **B**. The two forms of evolutionary change measured in this study. The trajectory change is a shift in which differentiation route an ortholog is expressed on; the temporal change is a shift in where along a shared route it is expressed. **C**. Single-cell RNA-seq datasets and developmental windows. Source datasets are identified by PMID. Shading indicates the neural production period. **D**. Trajectory inference. A scalar pseudotime gradient is converted into a per-cell velocity vector in the space of the first 50 principal components; a continuous vector field is reconstructed from these velocities; and the developmental path of highest probability between two cell states is recovered by least action path optimisation with expectation maximisation (Methods). **E**. Integrated datasets. UMAP embeddings coloured by consensus cell type (seven populations, annotated under a single marker-based scheme) and by developmental stage, with per-species embeddings below. **F**. The 13 inferred trajectories for human, projected onto the first three principal components; cells coloured by consensus cell type. **G**. Canonical fate markers along three representative trajectories in all four species. The shading marks the lineage in which each marker pair is expected. **H**. TrajMammal (**trajmammals.com**), a resource returning per-gene expression across all inferred trajectories in each species.

To infer developmental trajectories within these matched windows, we converted the gene expression gradient of each atlas into a per-cell velocity vector in the space of the first 50 principal components, reconstructed a continuous vector field from these velocities, and recovered the paths of highest probability by least action path optimization coupled with expectation maximization (**Figure 1D**, Methods). This approach captures the momentum of expression change across a cell population directly. Because trajectory comparison between species requires cells to carry consistent identities, we did not inherit the labels of the source atlases. Instead, all four datasets were re-annotated under a single marker-based scheme, which resolved each into seven populations: vRG, IPC-ExN, oRG, tRG, Glia-IPC, Astrocyte, and OPC-oligo, excluding terminal postmitotic neuronal subtypes for focusing on continuous differentiation trajectories of proliferative cell types (30,299, 51,324, 25,758, and 7,698 cells in human, macaque, mouse, and ferret, respectively; **Figure 1E**). The scheme recovered conserved marker expression in every species, although the proportion of each cell type varied between them (**Supplementary Figure S1**). Correspondence between the resulting cell types was near-complete, with 40 of 42 cross-species comparisons returning a two-way best match (95.2%, **Supplementary Figure S2**). These results demonstrate that the cell identities are defined equivalently in all four species.

This framework yielded a comparable set of developmental trajectories in each species. Projected onto the first three principal components of the human cells, the reconstructed vector field placed all 13 trajectories within the embedding (**Figure 1F**; other species in **Supplementary Figure S3**). Canonical fate markers showed the expected restriction along these routes in all four species, with *NEUROD1* and *NEUROG2* confined to the neuronal trajectories, *SLC1A3* and *CST3* to the astrocyte trajectories, and *OLIG2* and *PDGFRA* to the OPC-oligo trajectories (**Figure 1G**). The inferred trajectories therefore correspond to the developmental routes they were intended to capture. To make this substrate openly accessible, we deposited the complete per-gene output in TrajMammal (**trajmammals.com**), an interactive web-based database that returns the trajectory-specific expression, stage composition, and cell-type proportions for any queried ortholog in the four species (**Figure 1H**). Together, these results establish standardized trajectories that provide the common substrate on which orthologous genes can now be tested for evolutionary change.

### Most gene orthologs are used broadly and conservatively across species

To systematically quantify evolutionary conservation in gene expression across four mammalian species, we mapped shared, one-to-one orthologs onto our developmental trajectory framework. For each gene, we evaluated its expression distribution across different developmental trajectories and its progression dynamics along each path. This allowed us to define three core metrics (**Figure 2A-C**). “**Lineage broadness**”: the number of developmental trajectories a gene actively occupies in a given species, ranging from 1 for strictly lineage-restricted genes to 13 for ubiquitous genes. “**Allocation divergence**”: The difference in how two species distribute a gene’s expression among available trajectories, quantifying evolutionary shifts in trajectory usage. “**Progression divergence**”: The difference in a gene’s expression dynamics along the progression of a shared trajectory, quantifying evolutionary shifts in developmental progression. Both divergence metrics are calculated as Jensen–Shannon distances bounded between 0 (perfectly conserved) and 1 (highly divergent), enabling direct, standardized comparisons between any species pair (**Supplementary Table S1**).

**Figure 2.**
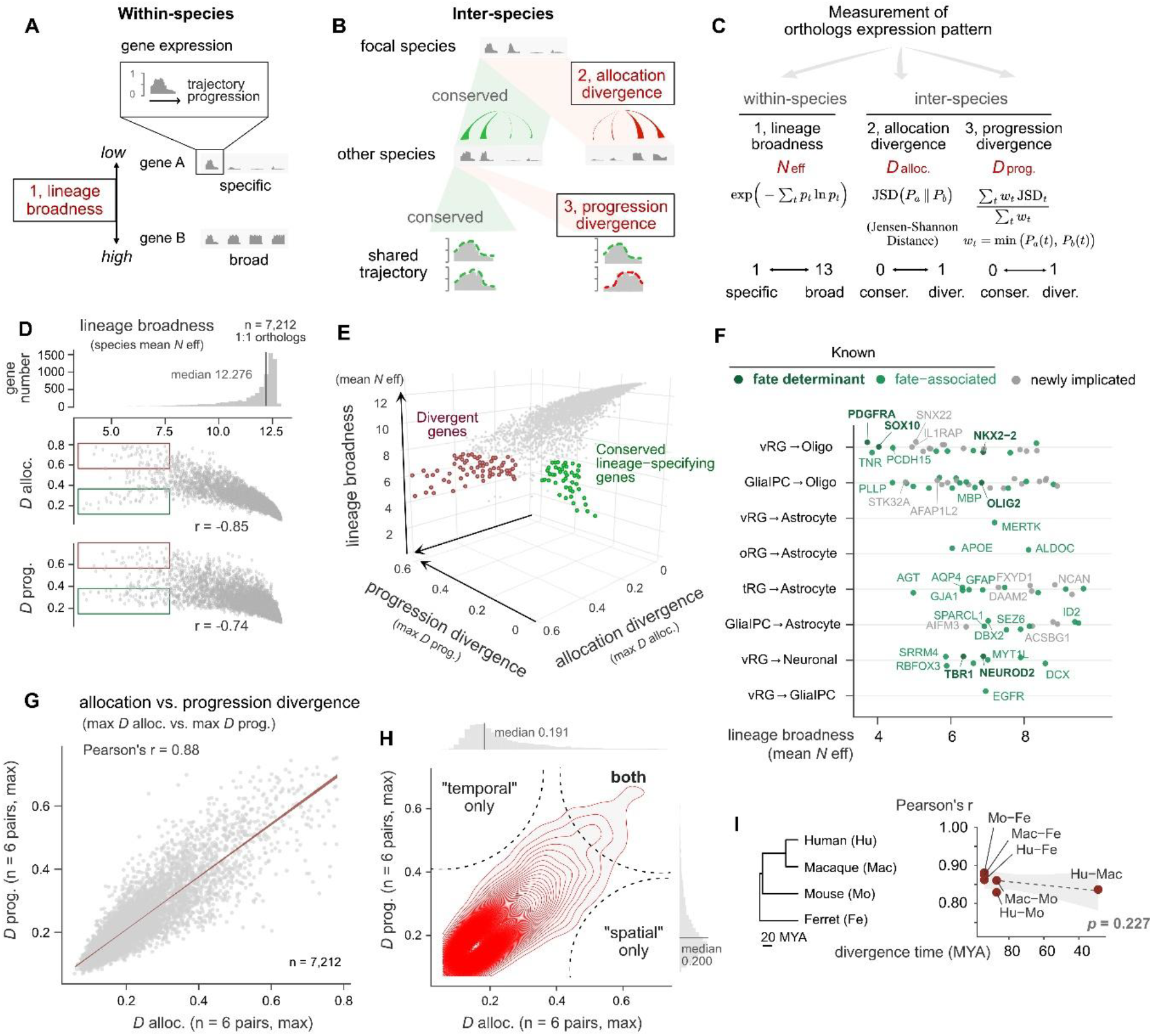
Genome-wide coupling of allocation and progression divergence. **A-C.** The three measures. Within a species, lineage broadness describes how many trajectories a gene occupies, from restricted (gene A) to broad (gene B). Between species, allocation divergence describes a change in which trajectories carry a gene, and progression divergence a change in its dynamics along a trajectory the species share. **C**. Definitions and ranges. Lineage broadness is the effective number of trajectories (***N*** eff), running from 1 to 13. Allocation divergence is the Jensen-Shannon distance between allocation distributions and progression divergence the mass-weighted mean of per-trajectory Jensen-Shannon distances, both bounded between 0 (conserved) and 1 (divergent). Weights ***w_t*** are the shared allocation mass of the two species on trajectory ***t*** (Methods). **D**. Lineage broadness across filtered orthologs (top, n = 7,212) and against each divergence metric (middle, bottom). Red boxes indicate narrowly deployed, highly divergent genes. Green boxes denote narrowly deployed but conserved genes on either axis. **E**. Combined plot of the three measures. Red points represent the highly divergent genes, corresponding to red boxes in (**D**). Green points mark “conserved lineage-specifying” genes, located in the lower corner of this space (see Methods). **F**. Conserved lineage-specifying genes placed on the trajectory carrying most of their expression, ordered by lineage broadness within each row. Colour indicates status: known fate determinant (dark green), known fate-associated gene (light green), or no reported role to our knowledge on that lineage (grey). **G-H**. Allocation against progression divergence for the retained orthologs, as a scatter with linear fit **(G)** and as a joint density with marginal distributions **(H)**. Dashed lines in **(H)** partition the space into regions in which one metric or both are elevated. **I**. Coupling strength against species splitting time. Each point is the correlation between allocation and progression divergence within one species pair, with the species tree at left. Dashed line, weighted regression on Fisher-transformed correlations. Correlations are Pearson unless stated; in **(I)**, confidence intervals are derived from the Fisher z transform and the regression is weighted by n − 3. Gene-level values for all metrics are given in **Supplementary Table S1**. Conserved lineage-specifying genes are listed in **Supplementary Table S2**.

Because single-cell expression metrics are highly sensitive to sampling noise at low transcript levels, we established a strict detection limit for minimal expression to be considered in further analysis (**Supplementary Figure S4A-C**). Across the 9,003 one-to-one orthologs, lineage broadness rose with expression magnitude, from a mean of approximately 2.5 effective trajectories among the weakest level genes to a plateau near 12 (**Supplementary Figure S4A**). Both divergences ran in the opposite direction, highest at low expression magnitude and lower at middle-to-high expression (**Supplementary Figure S4B**). To isolate true biological divergence from technical noise, we constructed an expression-matched null model by pairing randomly selected genes of similar abundance. Comparing our ortholog data against this null model revealed a clear threshold (maximum species expression of 0.023) where observed similarities exceeded chance (**Supplementary Figure S4C**). We restricted all subsequent analyses to the 7,212 orthologs that passed this robust expression threshold.

Applying these metrics genome-wide revealed a strikingly uniform transcriptional landscape: broad, stable expression is the default regulatory state across these four mammals. The vast majority of orthologs were expressed ubiquitously in multiple trajectories, with the median gene active in ∼12.3 of the 13 possible trajectories (**Figure 2D**). These broadly used genes also carried low divergence on both axes, and lineage broadness was negatively correlated with both allocation divergence (r = −0.85) and progression divergence (r = −0.74) (**Figure 2D**). Against this default background, a small set of genes departed from the relationship between broadness and divergence (green boxes in **Figure 2D**; conserved region in **Figure 2E**). These genes are highly exceptional, since the genes expressed in a smaller number trajectories tend to differ in their trajectory usage (red boxes in **Figure 2D**; divergent region in **Figure 2E**). The deep conservation of expression trajectories suggest their potential functional contribution of determining cell fates. To isolate them, we filtered for low allocation divergence, low progression divergence, and low lineage broadness, which returned 87 “conserved lineage-specifying” genes (**Figure 2F**; **Supplementary Table S2**; Methods). Mapping these 87 genes to their primary developmental trajectories successfully recovered canonical cell fate determinants (**Figure 2F**). *PDGFRA*^*45*^, *SOX10*^*46*^, and *NKX2*-2^47^ mapped to the oligodendrocyte routes, *OLIG2*^*48*^ to the Glia-IPC route toward oligodendrocytes, *TBR1*^*49*^ and *NEUROD2*^*50*^ to the neuronal route, and *AQP4*^*51*^, *GFAP*^*52*^, and *AGT*^*53*^ to the astrocyte routes. The filter uses no information about gene function, so its recovery of these genes in their correct trajectories indicates that both the inferred trajectories and the three measures behave as intended. Notably, 42% of this set have no reported role in their resident lineage, among them the poorly characterized kinase *STK32A* and the sorting nexin *SNX22* on the oligodendrocyte trajectories and the ion transport modulator *FXYD1* on the astrocyte ones. Given the accuracy with which the annotated members were placed, we speculate that these uncharacterized genes are candidate evolutionarily conserved regulators of the cell fate differentiation in which they reside.

### Coupled evolutionary shifts in allocation and progression

On top of this conserved background, we next examined the minority of gene orthologs that do diverge (red region in **Figure 2E**), and examine how the two forms of change, allocation and progression, are associated. Unexpectedly, we found a tight correlation of the two divergences across the 7,212 orthologs (Pearson r = 0.88; **Figure 2G**). The joint distribution of these metrics revealed a dense cluster of highly conserved genes that transitions into a distinct ridge along the diagonal (**Figure 2H**). Conversely, the off-diagonal regions were notably sparse: very few genes altered their trajectories while maintaining their progression dynamics, or vice versa. Therefore, when an ortholog undergoes evolutionary divergence, shifts in its trajectory allocation tend to coincide with shifts in its developmental progression.

To ensure this strong correlation reflects biological coupling rather than a technical artifact, we performed a series of rigorous mathematical controls. By default, progression divergence is weighted by the shared expression of a species pair within a given trajectory, which inherently links it to allocation. However, calculating an unweighted average across trajectories yielded a nearly identical correlation (r = 0.82; **Supplementary Figure S4D, E**), as did applying a fixed weight independent of the species pair (r = 0.85; **Supplementary Figure S4G, H**). The strong correlation also persisted after mathematically regressing out expression magnitude (r = 0.86; **Supplementary Figure S4I, J**) and remained robust when analyzing genes within narrow expression bins. Furthermore, this coupling was observed consistently across all six species pairs, showing only a slight, non-significant decline toward more recent evolutionary splits (*p* = 0.227; **Figure 2I**). Collectively, these mathematical controls validate this coupling as a genuine biological phenomenon. It reveals that while broad, conserved expression is the default state across the genome, genes that break from this constraint tend to change in allocation and in progression at once.

### Individual genes carry coupled changes on the human phylogenetic branch

Our genome-wide analysis revealed a robust correlation between allocation and progression divergence, yet its reliance on aggregated pairwise comparisons presents a critical limitation. A gene labelled as divergent could have accumulated its allocation change and its progression change on different phylogenetic branches, tens of millions of years apart. To resolve this and pinpoint regulatory modifications that evolved exclusively within the human branch, we developed a branch-resolved “contrast” metric (**Figure 3A–C**). For each gene, we calculated the expression divergence of humans from two non-primate outgroups (mouse and ferret), and subsequently subtracted the corresponding divergence of the macaque. This subtraction systematically filters out shared primate traits and macaque-specific adaptations, isolating only the evolutionary divergence that is specific to human species. Genes yielding the highest positive contrast scores were designated as having an “accelerated” human branch (**Figure 3C**). By applying this targeted metric to our two axes of divergence, we defined a core set of individual genes that underwent strictly human-biased “reallocation” (shifts in trajectory allocation) and human-biased “retiming” (shifts in developmental progression) (**Figure 3B, D, E**).

**Figure 3.**
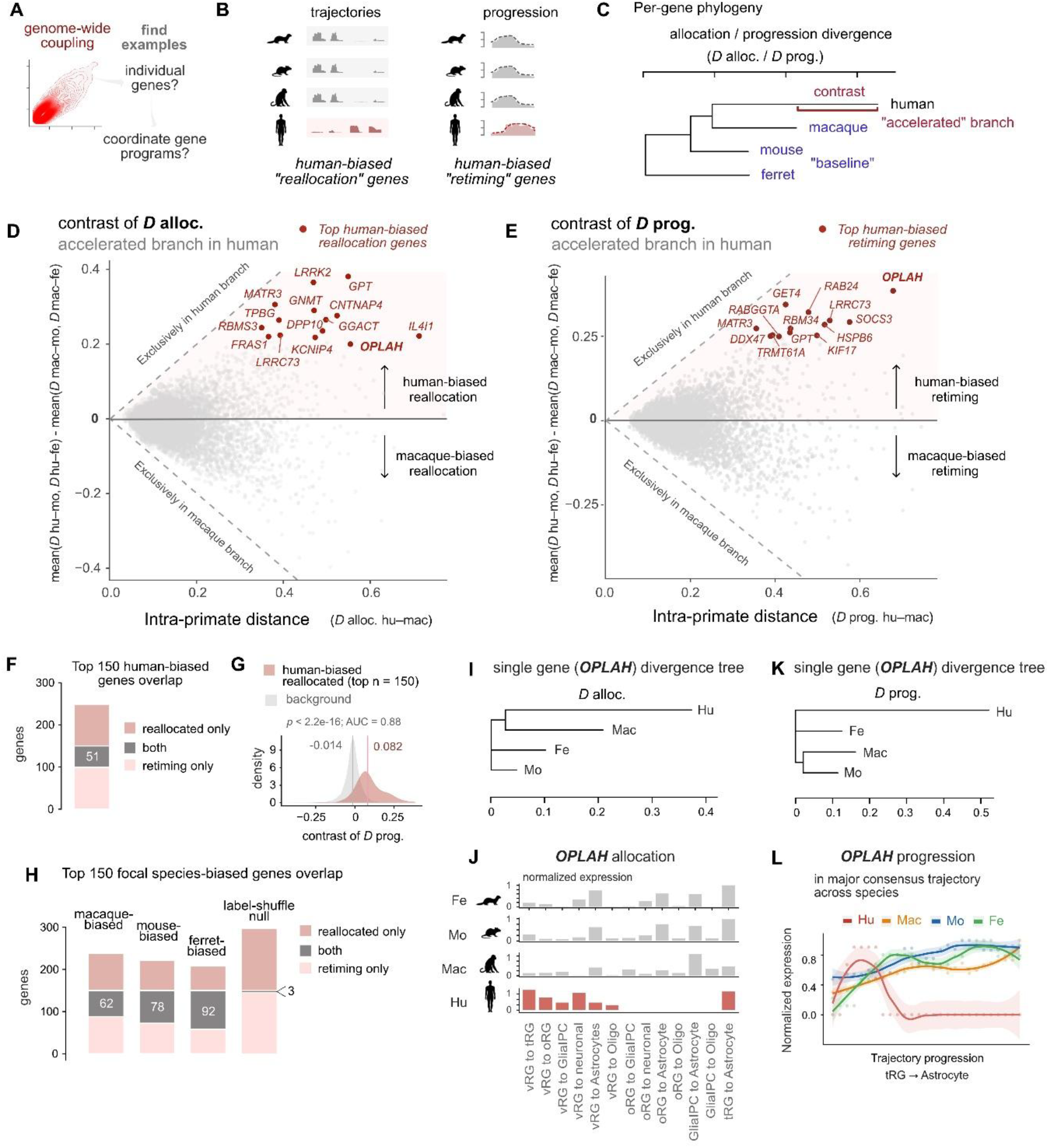
A branch-resolved method isolates coupled divergence on the human branch. **A.** Genome-wide coupling raises two questions, whether individual genes carry both changes in the same phylogenetic branch and whether such genes form coordinated programs. **B**. The two gene classes sought. Human-biased reallocation genes shift which trajectories carry their expression while the other species remain alike; human-biased retiming genes shift their dynamics along a shared trajectory. **C**. Assigning divergence to a branch. Genes with positive contrast values are described as having an “accelerated” human branch. **D-E**. Lineage contrast against intra-primate distance, computed on allocation divergence **(D)** and progression divergence **(E)**, for the 7,212 retained orthologs. Points above zero indicate human-biased change and below zero macaque-biased change; diagonals bound the region in which divergence is attributable entirely to one branch. Red points mark the top genes of each contrast. **F**. Overlap between the top human-biased reallocation and retiming gene sets (n = 150 each). **G**. Progression contrast of the top human-biased reallocation genes (n = 150) against all remaining orthologs (n = 7,062). Vertical lines mark medians (0.082 and −0.014). The reallocation set is shifted toward human-accelerated progression (one-sided Mann-Whitney *p* < 2.2 × 10^−16^, AUC = 0.88). **H**. The same overlap computed with each species in turn as the focal branch, together with a null in which gene labels on the progression contrast were shuffled before selection (mean of 1,000 permutations per focal species). **(I, K)** Per-gene divergence trees for *OPLAH*, built from pairwise allocation divergence **(I)** and progression divergence **(K)**. Scale bars in divergence units. **J**. *OPLAH* allocation across the 13 trajectories in each species, normalised within species. **L**. *OPLAH* progression along tRG to astrocyte, the trajectory carrying most of its expression in all four species. Points are per-step values and lines are LOESS fits with 95% confidence bands. Remaining trajectories in **Supplementary Figure S5**. Trees in **(I, K)** are *BIONJ* fits to the six pairwise divergences of a single gene; no bootstrap support is available for a per-gene tree. Densities in **(G)** are kernel estimates over all genes in each set. Top gene sets for every focal species are listed in **Supplementary Table S3**.

Both contrast metrics revealed clear statistical outliers against a highly conserved baseline centered near zero (**Figure 3D, E**). The top human-biased reallocation genes included *LRRK2, GPT, MATR3*, and *CNTNAP4*, while the top retiming genes included *OPLAH, SOCS3, RAB24*, and *LRRC73* (**Figure 3D, E**). Directly comparing these two rankings confirmed our previous genome-wide observations at the single-gene level: of the top 150 genes in each category, 51 were shared (including *OPLAH, GPT*, and *MATR3*; **Figure 3F**). Furthermore, genes selected solely for a significant shift in trajectory allocation simultaneously exhibited a marked shift in developmental progression (median retiming contrast 0.082 versus −0.014 for background; one-sided Mann–Whitney *p* < 2.2 × 10^−16^, AUC = 0.88; **Figure 3G**). These results demonstrate that the coupling observed genome-wide also holds within individual genes and on a single branch. Furthermore, this coupled divergence is not a uniquely human trait. Repeating the contrast procedure using macaque, mouse, or ferret as the focal species yielded overlaps of 62, 78, and 92 genes, respectively, compared to a label-shuffled null expectation of approximately 3 (**Figure 3H, Supplementary Table S3**). The overlap increased with the evolutionary depth of the branch tested, consistent with the mild strengthening of coupling toward deeper splits observed genome-wide (**Figure 2I**).

To visualize this coupled divergence at the single-gene level, we examined *OPLAH*, 5-oxoprolinase, the top-ranked gene in both the allocation and progression contrasts (**Figure 3D, E**). In macaque, mouse, and ferret, *OPLAH* expression is strictly confined to terminal astrocyte lineages and upregulates late in development. In stark contrast, human *OPLAH* expression has dramatically expanded across vRG-rooted trajectories (reallocation; **Figure 3J**) and temporally shifted to peak early in development before declining (retiming; **Figure 3L**; remaining trajectories in **Supplementary Figure S5**). Per-gene divergence trees placed both changes exclusively on the human branch (**Figure 3I, K**). Together, these results demonstrate that coupled allocation and progression divergence indeed arises simultaneously on a single phylogenetic branch, validating our contrast metric as a robust tool for discovering human-specific regulatory shifts.

### A subset of synaptic genes are reallocated and retimed on human oligodendrocyte lineage

Because evolutionary shifts frequently rewire entire molecular pathways rather than single targets, we next asked whether these human-biased genes evolve independently or converge into shared regulatory networks. We clustered them by their allocation distributions using consensus Weighted Gene Co-expression Network Analysis (WGCNA; **Figure 4A, B**). This procedure yielded four clusters that together captured 78% of the genes (**Figure 4C**). In humans, each gene cluster exhibited relatively strict lineage specificity, whereas the orthologous genes in the macaque, mouse, and ferret maintained broad, ubiquitous expression (**Figure 4D–G**). Human genes in these clusters occupied 7.4 to 9.1 effective trajectories, compared with 10.3 to 11.4 in the other three species (**Figure 4H**). Cluster 1 (n = 38) was restricted to oRG and neuronal routes, Cluster 2 (n = 29) to oligodendrocyte routes, Cluster 3 (n = 29) to astrocyte routes, and Cluster 4 (n = 21) to tRG routes. These results demonstrate that the human-biased genes are not isolated units but form coherent programs, each confined in human to a distinct part of the developmental landscape.

**Figure 4.**
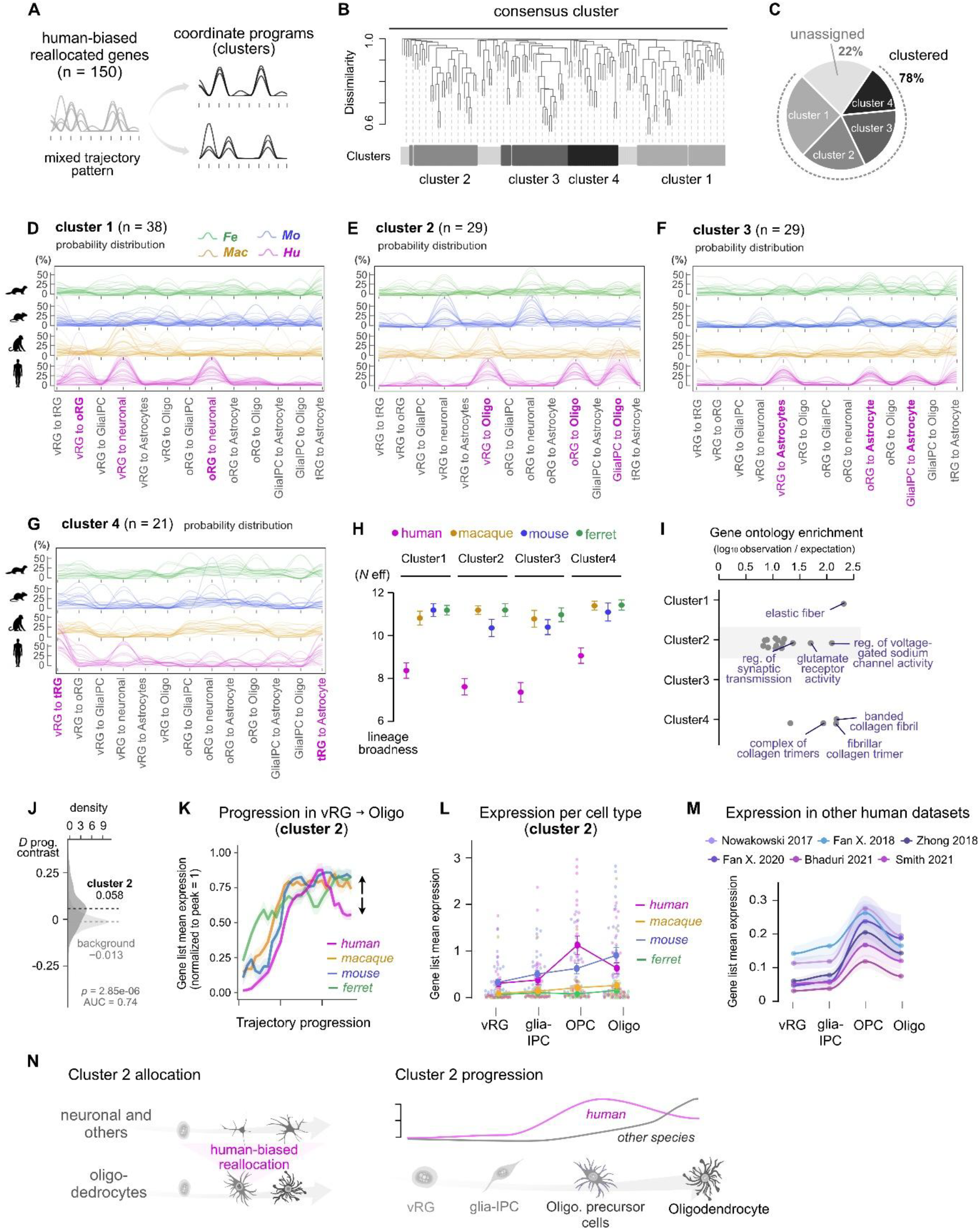
A synaptic gene network is deployed in the human oligodendrocyte lineage and peaks transiently in precursors. **A.** The top human-biased reallocation genes (n = 150) were clustered on their allocation distributions to ask whether they form coordinated programs. **B-C**. Consensus clustering of the 150 genes on their allocation across the 13 trajectories in all four species (52 features per gene; Methods). Dendrogram with cluster assignments **(B)** and the proportion of genes assigned to each cluster **(C)**; 78% were assigned and 22% left unclustered. **D-G**. Allocation distributions of each cluster in each species, shown as probability distributions. **H**. Lineage broadness of each cluster in each species. Human genes occupy an effective 7.4 to 9.1 trajectories against 10.3 to 11.4 in the other three species. **I**. Gene ontology enrichment for each cluster, shown as observation over expectation. Cluster 2 returned 14 significant terms (FDR < 0.05) against 1, 0 and 4 for clusters 1, 3 and 4; selected terms are labelled. **J**. Progression contrast of cluster 2 genes against all remaining orthologs. Dashed lines mark medians. One-sided Mann-Whitney *p* = 2.85 × 10^−6^, AUC = 0.74, for cluster 2 (n = 29) against background (n = 7,183). **K**. Cluster 2 mean expression along the vRG to oligodendrocyte trajectory in each species, normalised to the peak within species. Arrow marks the divergence in the second half of the trajectory, where human declines while the other species remain high. **L**. Cluster 2 mean expression across annotated cell types of the oligodendrocyte lineage in each species. Neuronal cell types in **Supplementary Figure S6. M**. The same cell-type profile in six independently re-annotated human datasets, each restricted to the matching developmental window (Methods). **N**. Summary. In human, cluster 2 is reallocated from neuronal and other routes onto the oligodendrocyte lineage, and within that lineage its expression is advanced to the precursor state. Data in **(H, L, M)** are mean ± SEM. Cluster membership and gene ontology results are given in **Supplementary Table S4**.

To uncover the biological functions driven by these human-specific regulatory networks, we performed Gene Ontology (GO) enrichment analysis on each cluster (**Figure 4I, Supplementary Table S4**). Cluster 2 was the only cluster with substantial enrichment, returning 14 terms (FDR < 0.05) centered on synaptic transmission and membrane excitability, including *GRIN2B, GRM7, KCNIP4, FGF12*, and *FGF14*. Across mammals, these genes are classically expressed in mature neurons; however, their robust shift into the oligodendrocyte trajectories was exclusively observed in humans. To rule out potential artifacts from our computational trajectory inference, we directly measured Cluster 2 expression across discrete, annotated cell types. Consistent with our trajectory mapping, expression of this cluster remained low across developing neuronal types in all four species, but was sharply elevated specifically within the human oligodendrocyte lineage (**Supplementary Figure S6**). These results reveal that a canonical synaptic gene program has been uniquely repurposed into the human oligodendrocyte lineage.

Given our previous finding that allocation and progression evolutionary shifts are tightly coupled, we hypothesized that the human-specific trajectory reallocation of Cluster 2 must be accompanied by a shift in developmental progression. Indeed, evaluating the progression contrast for Cluster 2 confirmed that this synaptic network is significantly retimed on the human branch (median 0.058 versus −0.013 for background; *p* = 2.85 × 10^−6^, AUC = 0.74; **Figure 4J**). Along the vRG-to-oligodendrocyte trajectory, the macaque, mouse, and ferret monotonically upregulated Cluster 2 late in development. In striking contrast, human expression of Cluster 2 peaked near the trajectory’s midpoint before declining steeply (**Figure 4K**). By anchoring this progression midpoint to discrete cell states, we found that human expression peaks transiently in oligodendrocyte precursor cells (OPCs) and drops in mature oligodendrocytes, whereas expression in the other three species peaks during maturation (**Figure 4L**). We successfully validated this transient OPC peak across six independent human transcriptomic datasets spanning the matching developmental window^54,55,56,57,58,59^ (**Figure 4M**, Methods), ensuring this signature is a robust biological phenomenon rather than an atlas-specific artifact.

Together, these findings demonstrate that Cluster 2 constitutes a synaptic program that, in human alone, is deployed in the oligodendrocyte lineage and concentrated in the precursor state rather than in mature oligodendrocytes (**Figure 4N**). This confirms that the coupled evolution of allocation and progression governs not just individual genes, but entire functional gene networks. Its immediate implication concerns the cells themselves: human OPCs are predicted to carry synaptic machinery transiently during development.

### Human-specific co-option of synaptic machinery in early oligodendrocyte precursor cells

To experimentally validate the presence of synaptic proteins in human OPCs and test for human-specificity, we examined cortical organoids derived from human, chimpanzee, and gorilla pluripotent stem cells (**Figure 5A**). We quantified the protein products of the NMDA receptor subunit GluN2B (*GRIN2B*), a representative Cluster 2 member, and the canonical postsynaptic scaffold PSD-95 (*DLG4*) in PDGFRα and SOX10 double-positive OPCs (**Figure 5B-D, Supplementary Figure S8**). PSD-95 is not itself a Cluster 2 gene; we included it because Cluster 2 is enriched for postsynaptic gene ontology terms (**Supplementary Table S4**), which suggests that these cells assemble a postsynaptic scaffold rather than expressing receptor subunits alone. By around two months of culture (DIV 60), both proteins were present in human OPCs. Volumetric reconstructions revealed discrete synaptic-like puncta within the PDGFRα- positive cellular volume, with GluN2B partially localized to the cell surface and PSD-95 concentrated intracellularly beneath the plasma membrane labeled by PDGFRα (**Figure 5C, D**). We quantified this signal and found it specifically elevated in early human cells. Human OPCs at DIV60 carried significantly more of both proteins than chimpanzee or gorilla OPCs in both signal volume and puncta density (**Figure 5F, G, I, J**; *p* < 0.05 for most comparisons; **Supplementary Table S5**). This expression is also highly stage-specific, with human OPCs carrying significantly more of both proteins at DIV60 than at DIV160 (*p* < 0.01; **Supplementary Table S5**).

**Figure 5.**
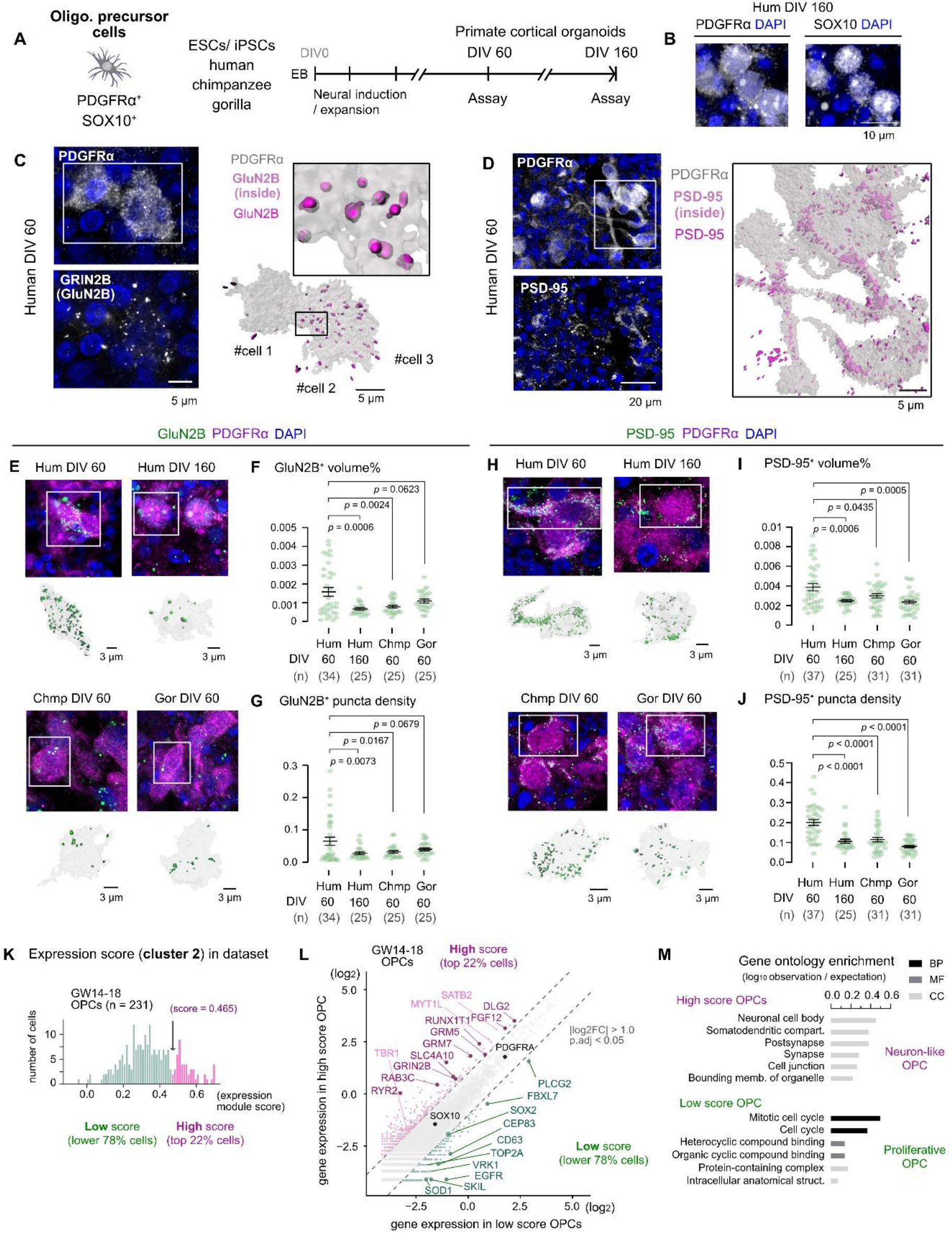
Human-specific co-option of postsynaptic machinery in early oligodendrocyte precursor cells. **A.** Cortical organoids were generated from human, chimpanzee and gorilla embryonic and induced pluripotent stem cells and assayed at Day in vitro (DIV) 60 and DIV160. Oligodendrocyte precursors were identified as PDGFRα and SOX10 double-positive cells in DIV160. **B**. PDGFRα and SOX10 immunostaining in human organoids at DIV160. **C-D**. Confocal imaging and volumetric reconstruction of GluN2B **(C)** and PSD-95 **(D)** within the PDGFRα-positive volume in human DIV60 organoids. GluN2B localises partly at the cell surface and PSD-95 predominantly beneath it. **E-G**. GluN2B in human OPCs at DIV60 and DIV160 and in chimpanzee and gorilla OPCs at DIV60. Representative images with reconstructions **(E)**, signal volume as a fraction of cell volume **(F)**, and puncta density **(G). H-J**. PSD-95 in the same conditions, shown as in **E-G. K**. Cluster 2 expression module score across human OPCs between GW14 and GW18 (n = 231 cells). The top 22% of cells (score above 0.465) were defined as high-score OPCs and the remainder as low-score OPCs. **L**. Gene expression in high-score against low-score OPCs. Coloured points are differentially expressed (|log2FC| > 1.0, adjusted *p* < 0.05); the OPC markers *PDGFRA* and *SOX10* are labelled in black and lie on the diagonal. Selected genes are labelled. **M**. Gene ontology enrichment for genes higher in each group, shown as observation over expectation and coloured by ontology domain. Full results in **Supplementary Table S7**. Data in **(F, G, I, J)** are mean ± SEM, with individual cells shown; GluN2b: n = 34, 25, 25, 25 and PSD-95: n = 37, 25, 31, 31 cells for human DIV60, human DIV160, chimpanzee DIV60 and gorilla DIV60, each from 2 independent organoid batches per species. Comparisons are Welch’s t-tests against human DIV60, with exact *p* values shown on each panel. Scale bars are given in each panel.

To determine whether this early developmental bias extends to the broader synaptic network *in vivo*, we tracked the expression trajectories of the complete Cluster 2 module alongside a comprehensive panel of 602 synaptic genes across chronological development—spanning from gestational week (GW) 14 to adulthood in humans, and embryonic day (E) 15 to postnatal day (P) 56 in mice (**Supplementary Figure S7A-D**). Human OPCs exhibited high prenatal transcription of this synaptic panel, which declined in later stages. In striking contrast, orthologous expression in mouse OPCs rose monotonically, only matching human transcript levels postnatally (**Supplementary Figure S7E-G**). Temporal clustering resolved this panel into sub-modules with distinct trajectories, and the sub-module containing *GRIN2B* (GluN2B) and *DLG4* (PSD-95) followed the same pattern as the organoid measurements: elevated early in human development and declining thereafter (**Supplementary Figure S7H-K, Supplementary Table S6**). Thus, the human-specific deployment of this synaptic network is largely transient and confined to the earliest phases of gliogenesis.

A closer inspection of the DIV60 organoids revealed that this human-specific protein expression is highly heterogeneous (**Figure 5C, D, E, H, Supplementary Figure S8**). A distinct subpopulation of human OPCs contained synaptic signal volumes several-fold higher than any hominid cells, while the remaining human OPCs expressed baseline levels comparable to non-human hominids (further examples in **Supplementary Figure S8**). To characterize this fraction, we scored early human OPCs *in vivo* on Cluster 2 expression (GW14 to GW18, n = 231 cells) and defined the top 22% as “high-score” OPCs (score > 0.465; **Figure 5K**). High-score and low-score cells expressed the core OPC markers *PDGFRA* and *SOX10* at comparable levels, confirming that the difference between them reflects heterogeneity within the OPC pool rather than lineage misidentification (**Figure 5L**). Differential expression analysis revealed that high-score OPCs were strongly enriched for additional synaptic genes (*DLG2, GRM5, SLC4A10*) as well as transcription factors known to regulate the fate of postmitotic neuron subtypes (*TBR1, SATB2, MYT1L*) (**Figure 5L**). Conversely, low-score OPCs were enriched for cell cycle and stem cell programs, including *EGFR, TOP2A*, and *SOX2*. Gene Ontology enrichment mirrored this divide, returning “postsynaptic” and “somatodendritic” terms for high-score cells versus “mitotic cell cycle” terms for low-score cells (**Figure 5M**; **Supplementary Table S7**). Therefore, the Cluster 2 network effectively segregates early developing human OPCs along an axis, from standard, highly proliferative glial precursors to a specialized “neuron-like” OPC state.

In summary, the human-specific co-option of a subset of postsynaptic genes from neuronal to oligodendrocyte trajectories exemplifies the tight coupling between allocation and progression divergences. Beyond transcriptomic changes, experimental evidence demonstrated a human-specific enrichment of postsynapse-like punctate structures marked by NMDA receptor and postsynaptic scaffold proteins at the midpoint of oligodendrocyte differentiation.

### Two types of OPCs occupy the same niche but differ in their molecular interface with outer radial glia

Both single cell transcriptomics and immunohistochemistry commonly suggest a heterogeneity of OPC population in terms of the expression of cluster 2 postsynaptic genes. To correlate these OPC subgroups and potential developmental functions, we reanalyzed spatial transcriptomic (MERFISH) atlases of the developing human cortex^28,60^. These datasets collectively quantify 300 targeted transcripts across 28 cortical sections, spanning from gestational week (GW) 15 through eight months of postnatal life (**Figure 6A**). We scored all 145,407 captured OPCs for expression of the Cluster 2 synaptic network, categorizing the top 20% as “neuronal-OPC” and the remainder as “proliferative-OPC” (**Figure 6B**).

**Figure 6.**
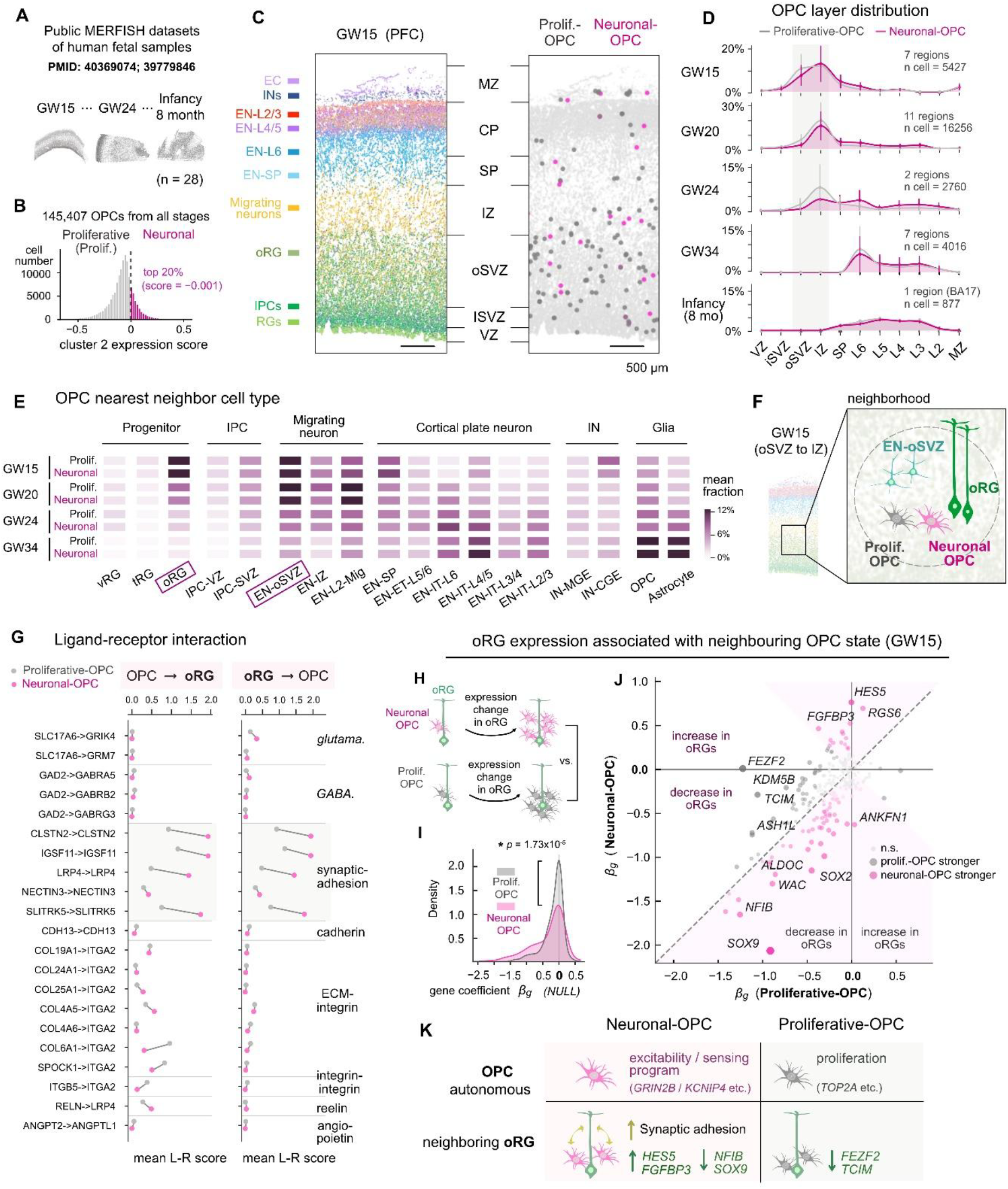
Neuron-like and proliferative OPCs occupy the same niche but differ in their molecular interface with outer radial glia. **A.** Published MERFISH datasets of human fetal cortex spanning GW15 to eight months postnatal (n = 28 samples). **B**. Cluster 2 expression score across all 145,407 OPCs. The top 20% of cells were defined as neuronal-OPC and the remainder as proliferative-OPC (or prolif. OPC). **C**. GW15 prefrontal cortex, coloured by cell type (left) and showing neuronal-OPC and proliferative-OPC cells in their laminar positions (right). **D**. Laminar distribution of neuronal-OPC and proliferative-OPC cells across developmental stages, from ventricular zone to marginal zone. OPCs are concentrated in the oSVZ and IZ at GW15 and in the cortical plate by infancy. Shading marks the oSVZ to IZ. Additional GW15 samples and regions in **Supplementary Figure S9. E**. Cell-type composition of each OPC’s immediate neighbourhood, derived from a Delaunay graph of cell centroids. **F**. Schematic of the GW15 oSVZ to IZ neighbourhood. **G**. Ligand-receptor scores in neuronal-OPC and proliferative-OPC neighbourhoods, in both directions, grouped by interaction class. Four homophilic synaptic-adhesion pairs score higher in neuronal-OPC neighbourhoods at the OPC-oRG interface (*q* < 0.05, Mann-Whitney U). **H**. Model design. For each gene in oRG cells, expression was regressed on the local density of neuronal-OPC and of proliferative-OPC cells, adjusting for cortical depth, sample, and the density of all other major cell types, giving a coefficient β for each OPC state (Methods). **I**. Distribution of β across the transcriptome in oRG (n = 388,610 cells), under each OPC state. Coupling is systematically stronger for neuronal-OPC (Wasserstein distance 0.20, Kolmogorov-Smirnov *p* = 1.73 × 10^-5^). **J**. β under neuronal-OPC against β under proliferative-OPC, per gene. Coloured points are significantly more strongly coupled to one state; selected genes are labelled. **K**. Summary. Cluster 2 expression separates an excitability program from a proliferative one within OPCs, and the two states are accompanied by different transcriptional profiles in adjacent oRG. Ventricular zone (VZ), inner subventricular zone (iSVZ), outer subventricular zone (oSVZ), intermediate zone (IZ), subplate (SP), cortical Plate (CP), marginal Zone (MZ). Per-gene coefficients are given in **Supplementary Table S8**.

We first evaluated the broad spatial distribution of these populations. At GW15 in the prefrontal cortex, the global OPC pool was heavily concentrated in the outer subventricular zone (oSVZ) and the intermediate zone (IZ) (**Figure 6C**). As development progressed, this distribution shifted dramatically: the early germinal-zone localization gave way to an intermediate-zone preference by GW20, and by infancy, OPCs resided predominantly within the cortical plate (**Figure 6D**). Thus, early oSVZ residency defines a tightly restricted developmental niche that precisely aligns with the temporal window of peak synaptic gene expression. However, this specific niche does not segregate the two OPC states. Neuronal-OPC and proliferative-OPC were laminarly indistinguishable at GW15 (Wasserstein distance 0.031, Kolmogorov– Smirnov *p* = 0.3213). This lack of spatial segregation persisted across all subsequent stages, cortical regions, and additional GW15 samples, with only a marginal, non-significant shift toward the pial surface observed (**Figure 6D, Supplementary Figure S9**).

Because the neuron-like state is not defined by macroscopic laminar position, we hypothesized that the two OPC populations might assemble distinct local microenvironments. We constructed a Delaunay neighbor graph from cell centroids in each tissue section to comprehensively map the immediate cellular contacts of every OPC (Methods). At GW15, the overall OPC population preferentially contacted outer radial glia (oRG) and migrating excitatory neurons of the oSVZ (EN-oSVZ), followed by subplate neurons, migrating layer 2 neurons, and CGE-derived interneurons (**Figure 6E, F**). This specific microenvironment dissolved over developmental time; by GW34, OPCs were no longer enriched near these early progenitors, but instead intercalated among mature cortical plate neurons and glia. Notably, neuronal-OPC and proliferative-OPC cells shared comparable neighbourhoods at every stage. Therefore, macroscopic position and local cellular composition are indistinguishable between the two OPC states, which leaves the difference between them to be sought in how they interface with their shared neighbors.

To identify molecular channels that distinguish these two OPC microenvironments, we systematically quantified ligand-receptor signaling scores between neuronal-OPC versus proliferative-OPC cells and their adjacent neighbors (**Figure 6G**). Despite the limited detectability due to only 300 probes included in the dataset, at the physical interface between OPCs and oRGs, four homophilic synaptic-adhesion pairs (*CLSTN2, SLITRK5, IGSF11*, and *LRP4*) scored significantly higher in neuronal-OPC neighborhoods compared to proliferative-OPC neighborhoods (q < 0.05). Because these are homophilic, these adhesion molecules are heavily co-expressed by both the neuron-like OPC and the adjacent oRG. In contrast, at the OPC-to-EN-oSVZ interface, only *CLSTN2* reached significance (**Supplementary Figure S10**). No glutamatergic or GABAergic pair (e.g., *SLC17A6* to *GRIK4* and *GAD2* to *GABRB2*) differed between the two states of OPCs. These results suggest a functional interaction of neuronal-OPC and neighboring cells through synaptic adhesions, potentially leading to the reaction in oRG.

We next investigated whether this enhanced synaptic adhesion is coupled to distinct transcriptional states within the neighboring cells. We modeled gene expression in neighboring oRGs and EN-oSVZ neurons as a function of the local density of neuronal-OPC versus proliferative-OPC cells, controlling for cortical depth, sample identity, and the density of all other major cell types (Methods). This model outputs a coefficient isolating the unique transcriptional association of each OPC state with its neighbor (**Figure 6H**). In oRG (n = 388,610 cells), this transcriptional coupling was systematically stronger for neuronal-OPC than for proliferative-OPC density across the gene panel (Wasserstein distance 0.20, Kolmogorov– Smirnov *p* = 1.73 × 10^−5^; **Figure 6I**). The transcripts strongly coupled to neuronal-OPC density in adjacent oRGs were dominated by downregulated glial fate-commitment regulators, including the astrocytic factors *SOX9, NFIB*, and *ALDOC*. Concurrently, the radial glia maintenance factor *HES5* and the FGF signaling modulator *FGFBP3* were significantly upregulated (**Figure 6J**). The combination of elevated *HES5* and *FGFBP3* with suppressed gliogenic markers suggests that oRGs remain in a less committed, more progenitor-like state when adjacent to neuron-like OPCs, although *SOX2*, also associated with the undifferentiated state, moved in the opposite direction and argues against that interpretation. This transcriptional signature was highly robust: 16 of the 20 most strongly neuronal-OPC-coupled genes survived rigorous label shuffling within the OPC pool (*n_permutation* = 2,000), including all aforementioned markers (**Supplementary Table S8**). Conversely, in EN-oSVZ neurons (n = 441,877 cells), this coupling was vastly weaker and less polarized (Wasserstein distance 0.09), yielding a mixed signature of neuronal fate markers and neurite outgrowth genes (**Supplementary Figure S10**). Overall, transcriptional coupling was approximately twofold stronger at the oRG interface than at the EN-oSVZ interface (‖β‖2 ≈ 7 against ≈ 4 across both senders), identifying oRG as the primary cellular target sensitive to the neuronal-like OPC state.

Together, these spatial analyses define a narrow developmental window at GW15 in which neuron-like OPCs and their oRG neighbors assemble a distinct microenvironment governed by elevated synaptic-adhesion signaling (**Figure 6K**). Within this niche, the neuronal state of an OPC is strongly associated with a delayed, less committed transcriptional profile in physically adjacent oRG. While spatial transcriptomics alone cannot definitively resolve the directionality of this interaction, these data raise the compelling hypothesis that an early human OPC subpopulation do not passively occupy the oSVZ niche, but instead deploy co-opted synaptic adhesion machinery to actively modulate oRG program and influence human cortical development.

## Discussion

In this study, we decoded the evolutionary rewiring of the cerebral cortical development by aligning continuous single-cell differentiation trajectories across the human, rhesus macaque, mouse, and ferret (**Figure 1**). By scoring orthologous genes on both trajectory usage (allocation) and expression timing along differentiation trajectories (progression), we uncovered a fundamental principle of transcriptomic evolution. Although broad, conserved expression is the mammalian default, regulatory divergence strictly forces genes to shift their spatial and temporal coordinates in tandem (**Figure 2**). To isolate how this coupled evolution shaped the human brain, we applied a branch-resolved contrast metric, revealing that a canonical synaptic gene network was uniquely redeployed into early human oligodendrocyte precursor cells (OPCs) (**Figures 3 and 4**). Cortical organoids confirmed that this synaptic machinery is translated into robust protein expression at significantly higher levels in human OPCs compared to those of chimpanzees and gorillas. We found a significant heterogeneity of human OPCs; neuron-like OPC subpopulation is defined by a notable level of synaptic gene expressions and the rest, proliferative-OPC, demonstrating enriched expression of genes related to proliferation and stem cell regulation (**Figure 5**). Finally, spatial mapping localized these neuron-like OPCs to the outer subventricular zone during a narrow prenatal window, where they actively engage oRGs via elevated synaptic-adhesion signaling to potentially maintain a less committed progenitor niche (**Figure 6**). These findings demonstrate that the coupled spatiotemporal rewiring of conserved gene networks can establish entirely novel cellular microenvironments, providing a distinct molecular engine for human cortical evolution.

### Evolutionary novelty and mechanism of the synaptic program in human OPCs

Oligodendrocyte precursor cells (OPCs) are the only glial cell type known to form true synapses with mature neurons^61,62^. This synaptic connectivity allows them to sense neural activity, directly regulate their own proliferation, and dynamically modulate local circuits^63–68,69–71^. However, these functional insights derive almost entirely from mouse and zebrafish models during postnatal myelination. Recently, a distinct synaptic gene program was also reported in human fetal OPCs; immunopanned OPCs from the second-trimester cortex revealed a transcriptionally distinct, germinal zone-localized cluster enriched for synaptic transcripts^72^. Our evolutionary measurements agree with that description and further provide evidence of human-specificity from the comparative analyses. The program was absent from the macaque, mouse, and ferret trajectories, and the postsynaptic proteins were far less abundant in chimpanzee and gorilla organoids at matched stages (**Figures 4 and 5**). The synaptic state of early OPCs is therefore not a general property of mammalian or even of hominid development, but rather a unique regulatory divergence that evolved exclusively on the human phylogenetic branch.

The molecular mechanisms enabling a human glial precursor to activate a canonical neuronal gene network can be tested directly. In non-neuronal cells, neuronal transcripts are typically repressed by the REST complex, which binds to RE1 elements present in many Cluster 2 genes, including *GRIN2B*^*73–77*^. Releasing this repression is sufficient to switch these genes on. A human-specific reduction in REST occupancy at these elements, or sequence change within the elements themselves, would account for our observations without any change to protein coding sequence, and existing epigenomic datasets are sufficient to test it. Alternatively, the transcription factors expressed in these cells (*TBR1, SATB2*, and *MYT1L*) may drive synaptic gene expression directly **(Figure 5L)**^49,78–80^. The robust expression of *MYT1L*, classically defined as a repressor of non-neuronal fates^79^, is particularly striking. Its robust co-expression in a PDGFRα-positive, SOX10-positive cell suggests that the transcriptional boundary separating neuronal and glial lineages is notably more permissive in humans than the rodents. It is worth experimentally examining the functional roles of these specific transcription factors in specifying the “neuronal” state or type of human OPC population in cortical organoids of both human and non-human hominids.

### A transient synaptic state of human OPC with implications for glioma vulnerability

Determining whether this neuronal-like profile represents a permanent OPC sub-lineage or a transient, reversible developmental state is critical for interpreting its biological function. Multiple lines of evidence in our dataset strongly favor a transient state. Both neuron-like and standard proliferative OPCs occupied identical laminar positions within the fetal cortex at every developmental stage examined. Furthermore, they shared indistinguishable local cellular neighborhoods and expressed core glial lineage markers (*PDGFRA* and *SOX10*) at comparable levels (**Figures 5L and 6D, E**). If these populations represented permanently divergent subtypes, we would expect to observe clear spatial segregation or differential lineage marker expression. This interpretation aligns with established rodent literature, which demonstrates that while synaptic inputs to OPCs are maintained through cell division, they are rapidly dismantled the moment differentiation begins^81,82^. We therefore propose that early human OPCs temporarily adopt a synaptic state to actively engage their local cellular environment. Although our spatial transcriptomic data confirm the enrichment of synaptic adhesion molecules at the interface between neuron-like OPCs and neighboring outer radial glia (**Figure 6G**), these molecules could mediate the formation of bona fide electrochemical synapses or simply act as structural adhesive scaffolds. Definitively resolving these mechanistic hypotheses will require targeted functional assays. Live imaging of uniquely labeled OPCs expressing fluorescently tagged synaptic proteins in human cortical organoids would reveal whether individual cells dynamically enter and exit this synaptic state, directly distinguishing a transient state from a rigid subtype. Additionally, pairing electron microscopy with patch-clamp electrophysiology in fetal slices or organoids is required to determine whether these accumulated proteins assemble into electrophysiologically active synapses or function purely as adhesive tethers. Finally, employing retrograde synaptic tracing from neuron-like OPCs in human fetal tissue or organoids promises to map the exact presynaptic inputs driving this specialized glial population.

Beyond normal development, this transient synaptic capacity carries profound pathological implications for neuro-oncology. High-grade gliomas are known to form functional, AMPA receptor-dependent synapses with neurons, utilizing these neuronal inputs to directly drive tumor growth and invasion^83–85^. The synapse-forming compartment within these tumors specifically resembles an OPC-like state, and recent models of diffuse midline glioma predict GABAergic neuron-to-glioma synapses based on the premise that tumor cells hijack normal OPC biology^86^. If human OPCs possess an intrinsic, specialized synaptic state during fetal development that rodent OPCs lack, the developmental substrate that gliomas exploit may be far more abundant and hardwired in the human brain than animal models capture. It is highly plausible that the human-specific evolutionary adaptation described here, placing synaptic machinery in a proliferative germinal precursor, is the precise developmental vulnerability that these aggressive tumors hijack.

### Coordinated evolution of spatial and temporal gene expression and its mechanistic basis

The human OPC program changed in two respects at once. The synaptic genes moved onto a different trajectory, and they came to peak at a different point along it. Evolutionary biology has long had names for changes of these two kinds: a feature that arises from a different part of the developing embryo than in the ancestor is *heterotopic*, and one that arises at a different point in the developmental sequence is *heterochronic*^*87–89*^. Historically, distinguishing between heterotopy and heterochrony at the macroscopic morphological level has proven difficult, frequently leading to the controversial reassignment of specific evolutionary adaptations from one category to the other^90,91^. Our single-cell trajectory framework gives each a concrete definition at the single cell-level gene expression: allocation divergence for heterotopy, progression divergence for heterochrony. Measured this way, these two evolutionary axes are tightly coupled across the mammalian genome; genes that undergo evolutionary change in their trajectory allocation tend to also change in their developmental progression timing (**Figures 2 and 3**). This genome-wide co-occurrence raises a mechanistic question: what types of biological mechanisms drive this tight correlation? We consider two hypothetical models.

In the “one-step” model, changes along the two axes represent a single evolutionary event manifesting in two dimensions. This one-step evolution relies on a concrete mechanistic basis. Regulatory logic of gene expression during development routinely carries positional and temporal information together; therefore, a single alteration to a regulatory element shift both coordinates at once. Sonic hedgehog in the neural tube provides a clear example. Its concentration specifies position along the dorsoventral axis, while its duration specifies which targets are expressed, so a change in the source of the signal or in how long it persists alters where and when together^92–94^.

In the “two-step” model, the heterotopic and heterochronic changes are distinct events coupled sequentially. We hypothesize that each single gene has a certain range of spatiotemporal expression niche and natural selection does not tolerate its expression outside the cell types and developmental stages where it is permitted to function. This stringent selective pressure confines the vast majority of the transcriptome to a highly conserved baseline, anchoring most genes in the low-divergence region where both allocation and progression remain stable (**Figure 2F and G**). Once either expression domain or timing of a given gene is changed during evolution, its expression will disappear if its ectopic function is harmful. Otherwise, the gene has to subsequently shift along the second axis to secure a new, permissive spatiotemporal niche where it can function in a novel cellular context. Here, the dense diagonal ridge of conserved genes and the sparse off-axis regions (**Figure 2G**) reflect the transience and instability of intermediate, single-axis states. The pattern might equally be called “all-or-none”: a conserved default in which place and time are togetherly constrained, and a displaced state in which both have moved.

Our data lean slightly toward the two-step model: the overlap between reallocated and retimed genes grew gradually with evolutionary branch depth (**Figure 3H**), and genome-wide coupling strengthened slightly over time, though it did not reach statistical significance (**Figure 2J**). A definitive test, however, requires species that diverged much more recently than humans and macaques (separated by 29 million years). The hominids (human, chimpanzee, and gorilla), which diverged 6 to 13 million years ago, offer an ideal comparison. Although *in vivo* single-cell atlases for these species remain unavailable, cortical organoids and other stem cell models of embryogenesis or organogenesis serve as an effective proxy to address this question^95,96^. Under the two-step model, these recently diverged branches should harbor a substantial fraction of genes displaced on only one axis. Conversely, under the one-step model, even the most recent evolutionary changes would already fall along the coupled diagonal.

### Heterotopy-heterochrony coupling in other systems

If coupling is a general property of developmental gene expression, it should be observable in other well-characterized cases. Indeed, several existing studies of cortical development highlight examples where spatial and temporal shifts co-occur. For instance, in our recent study, rat cortical progenitors generate deep-layer neurons for a longer duration than those in mice by autonomously secreting and sustaining Wnt signaling. This shift is at least driven by the ectopic expression of *LMX1A*, which is an upstream transcriptional factor driving Wnt ligand expression in the signaling center cortical hem, within the progenitors themselves, rather than restricting it to the neighboring morphogen secreting center as in mice^27^.

Coupling has also been observed outside the cerebral cortex. In Darwin’s finches, differences in beak depth accompany *BMP4* expression that is both broader in domain and earlier in onset, while differences in beak length accompany calmodulin expression in a distinct domain^97,98^. In bats, forelimb elongation involves prolonged and spatially extended *BMP2* signaling in the digits. Furthermore, transferring the bat *Prx1* limb enhancer into mice lengthens the forelimb, demonstrating that a single *cis*- regulatory change can alter both expression domain and duration simultaneously^99,100^. In snakes, the loss of regional identity along the trunk involves both shifted *Hox* boundaries and an altered segmentation clock^101,102^. Although these well-characterized cases demonstrate that spatial and temporal shifts co-occur, they do not prove that this genome-wide coupling is a universal rule across other organs. Systematically testing this principle elsewhere requires time-resolved, cross-species single-cell atlases. While such resources remain limited or absent for most organs^24^, their construction is a critical imperative to elucidate how broadly this coupled evolutionary mechanism operates across diverse tissues.

More broadly, these findings carry implications for the adjustable potential (“Evolvability”) of gene expression changes. Rather than varying independently, gene expression appears to evolve along a constrained, coupled diagonal within spatiotemporal space. Although this restricts available evolutionary trajectories, it simultaneously catalyzes novelty: relocating a gene program to a new cell lineage inherently alters its developmental timing, forcing it into a novel context with different molecular partners. Human-specific co-option of a synaptic gene network to an OPC subpopulation and its potential impact on oRG program may provide an example. Whether this mechanism fully explains how a largely shared mammalian genome builds organs of vastly different sizes and functions, typically exemplified by the cerebral cortical neural circuit, remains an open question. Nevertheless, this coupling reveals a highly efficient mechanism for rapid adaptation to distinct ecological demands, firmly establishing the spatiotemporal redeployment of existing genetic programs as a primary driver of evolutionary innovation.

## Resource availability

### Lead contact

Requests for further information and resources should be directed to and will be fulfilled by the lead contact, Ikuo K. Suzuki.

### Materials availability

This study did not generate new unique reagents. Primate pluripotent stem cell lines details are listed with their identifiers in the key resources table. Distribution of the non-human hominid lines is subject to the terms of the originating material transfer agreements; requests should be directed to the lead contact.

### Data and code availability

Inferred cell trajectories from each species, cell–cell transition probability matrices, and divergence metrics generated in this study are publicly available at Zenodo (https://zenodo.org/records/22026149. DOI: 10.5281/zenodo.22026149). Trajectory-resolved expression for any queried ortholog can additionally be browsed interactively at TrajMammal (https://trajmammals.com).

Raw and reconstructed imaging data for the cortical organoid experiments are publicly available at the BioImage Archive (biostudies; accession: S-BIAD3996). This record contains the unmodified confocal z-stacks, the corresponding Imaris reconstruction files, and the per-cell signal volume and puncta density values used for quantification.

Published single-cell transcriptomic atlases reanalyzed in this study were generated by others and are available at their original repositories, as listed below.

Human developing cortex scRNA-sequencing dataset: Wang et al.^28^ (https://datadryad.org/dataset/doi:10.5061/dryad.2280gb612); Fan et al.^56^ (GEO GSE120046, https://www.ncbi.nlm.nih.gov/geo/query/acc.cgi?acc=GSE120046); Nowakowski et al.^57^ (UCSC cell browser, https://cortex-dev.cells.ucsc.edu); Bhaduri et al.^54^ (UCSC cell browser, https://cells.ucsc.edu/?ds=dev-brain-regions); Fan et al.^55^(GEO GSE103723, https://www.ncbi.nlm.nih.gov/geo/query/acc.cgi?acc=GSE103723); Zhong et al.^59^ (GEO GSE104276, https://www.ncbi.nlm.nih.gov/geo/query/acc.cgi?acc=GSE104276); Smith et al.^58^ (Figshare dataset: https://figshare.com/s/64b648891e4817efb123).

Rhesus macaque developing cortex: Micali^29^ (GEO GSE226451, https://www.ncbi.nlm.nih.gov/geo/query/acc.cgi?acc=GSE226451).

Mouse developing cortex: Daniela J. Di Bella^30^ (GEO GSE153164, https://www.ncbi.nlm.nih.gov/geo/query/acc.cgi?acc=GSE153164); Gao et al.^31^ (https://allen-developmental-mouse-atlas.s3.amazonaws.com/scRNA/DevVIS_scRNA_processed.h5ad).

Ferret developing cortex, Lucia Del-Valle-Anton et al.^33^ (GEO GSE234305, https://www.ncbi.nlm.nih.gov/geo/query/acc.cgi?acc=GSE234305); Merve Bilgic ^32^(DDBJ, https://ddbj.nig.ac.jp/public/ddbj_database/dra/fastq/DRA016/DRA016867/).

Published spatial transcriptomic (MERFISH) atlases of the developing human cortex reanalyzed in this study are available at Wang et al.^28^ (https://datadryad.org/dataset/doi:10.5061/dryad.2280gb612) and Qian et al.^60^ (https://zenodo.org/records/14422018).

All original code used for trajectory reconstruction, divergence metric calculation, branch-resolved contrast analysis, gene clustering, and spatial neighbourhood modelling is publicly available at https://github.com/XD-Sheu/Coupling-divergence_2026 and is archived at Zenodo (DOI: 10.5281/zenodo.22026149).

## Acknowledgments

This research was supported by AMED (grants JP25gm7010016, JP24tm0524007, and JP20gm6310006), JST FOREST (JPMJFR214T), and MEXT KAKENHI (JP22H02628, JP20H04860, JP25K02280, and JP26H01794). We also gratefully acknowledge support from the Molecular Biology Society of Japan (MBSJ) via the Tomizawa Jun-ichi & Keiko Fund for Young Scientists, the Takeda Science Foundation, and the SECOM Science and Technology Foundation. We acknowledge the authors of the published single-cell and spatial transcriptomic atlases reanalyzed in this study for making their data openly available.

## Author contributions

I.K.S. and X.D.S. conceptualized the study. X.D.S., R.A., Y.N., J.Y., and I.K.S. developed the computational methods. X.D.S. performed the computational analyses. Y.Y.Y. and X.D.S. performed the organoid and imaging experiments. X.D.S. and I.K.S. wrote the paper. I.K.S. supervised the research and acquired funding.

## Declaration of generative AI and AI-assisted technologies in the writing process

During the preparation of this work, the authors used Gemini and Claude to improve readability and perform grammar checks. After using these tools, the authors reviewed and edited the content as needed and took full responsibility for the content of the published article.

## Methods and Materials

### Experimental model and study participant details

Primate cortical organoids were generated from four pluripotent stem cell lines: human WA09 (H9) hESCs^123^ (female), chimpanzee MARI iPSCs^103^ (female) and IKU iPSCs^120^ (female), and gorilla WER1 iPSCs (courtesy from Masanori Imamura laboratory, Kyoto University), following the protocol described in detail in the method details. Line names, sources, and identifiers are listed in the key resources table. No non-human primates were handled and no animal experiments were performed in the course of this study. No new sequencing data were generated in this study. All single-cell and spatial transcriptomic datasets analyzed here were generated by others and obtained from public repositories; sources, species, developmental stages, and accessions are listed in the key resources table and in the data and code availability section. Study participant details for the human datasets are as reported in the original publications.

### Key resources table

| REAGENT or RESOURCE | SOURCE | IDENTIFIER |
| --- | --- | --- |
| **Antibodies** |  |  |
| Goat polyclonal anti-PDGFRα | R&D Systems | Cat#AF1062; RRID: AB_2236897 |
| Mouse monoclonal anti-SOX10 (clone 1074) | abcam | Cat# ab216020; RRID:AB_2847913 |
| Rabbit polyclonal anti-PSD95 | proteintech | Cat# 20665-1-AP; RRID: AB_2687961 |
| Rabbit polyclonal anti-GRIN2B | proteintech | Cat# 21920-1-AP; RRID: AB_11232223 |
| Alexa Fluor 488 donkey anti-rabbit IgG (H+L) | Abcam | Cat# ab150073; RRID:AB_2636877 |
| Alexa Fluor 555 donkey anti-mouse IgG (H+L) | Abcam | Cat# ab150106; RRID:AB_2857373 |
| Alexa Fluor 647 donkey anti-goat IgG (H+L) | Abcam | Cat# ab150131; RRID:AB_2732857 |
| **Chemicals, peptides, and recombinant proteins** |  |  |
| Matrigel, growth factor reduced | Corning | Cat# 354277 |
| Y-27632 (ROCK inhibitor) | STEMCELL technologies | Catalog #72302 |
| Accutase enzyme solution | Sigma-Aldrich | Catalog #A696 |
| StemFit AK02N | stemfit | Catalog #RCAK02N |
| **Critical commercial assays** |  |  |
| STEMdiff™ Cerebral Organoid Kit | STEMCELL Technologies | Catalog #08570 |
| STEMdiff™ Cerebral Organoid Maturation Kit | STEMCELL Technologies | Catalog # 08571 |
| **Deposited data** |  |  |
| Inferred cell trajectories and cell–cell transition probability matrices | This paper | Zenodo:<br>https://doi.org/10.5281/zenodo.22026149 |
| per-ortholog expression divergence metrics across species | This paper | Zenodo:<br>https://doi.org/10.5281/zenodo.22026149 |
| Raw confocal images and Imaris reconstructions of oligodendrocyte precursor cells | This paper | BioImage Archive: S-BIAD3996 |
| Human, Rhesus macaque, Mouse and Ferret integrated scRNA-seq object | This paper | Zenodo:<br>https://doi.org/10.5281/zenodo.22026149 |
| Human developing cortex scRNA-seq and MERFISH | Wang et al.<sup>28</sup> | https://datadryad.org/dataset/doi:10.5061/dryad.2280gb612 |
| Human developing cortex MERFISH | Qian et al.<sup>60</sup> | https://zenodo.org/records/14422018 |
| Human developing cortex scRNA-seq | Fan et al.<sup>56</sup> | GEO GSE120046,<br>https://www.ncbi.nlm.nih.gov/geo/query/acc.cgi?acc=GSE120046 |
| Human developing cortex scRNA-seq | Nowakowski et al.<sup>57</sup> | UCSC cell browser, https://cortex-dev.cells.ucsc.edu |
| Human developing cortex scRNA-seq | Bhaduri et al.<sup>54</sup> | UCSC cell browser,<br>https://cells.ucsc.edu/?ds=dev-brain-regions |
| Human developing cortex scRNA-seq | Fan et al.<sup>55</sup> | GEO GSE103723,<br>https://www.ncbi.nlm.nih.gov/geo/query/acc.cgi?acc=GSE103723 |
| Human developing cortex scRNA-seq | Zhong et al.<sup>59</sup> | GEO GSE104276,<br>https://www.ncbi.nlm.nih.gov/geo/query/acc.cgi?acc=GSE104276 |
| Human developing cortex scRNA-seq | Smith et al.<sup>58</sup> | Figshare dataset:<br>https://figshare.com/s/64b648891e4817efb123 |
| Rhesus macaque developing cortex scRNA-seq | Nicola Micali<sup>29</sup> | GEO GSE226451,<br>https://www.ncbi.nlm.nih.gov/geo/query/acc.cgi?acc=GSE226451 |
| Mouse developing cortex scRNA-seq | Daniela J. Di Bella<sup>30</sup> | GEO GSE153164,<br>https://www.ncbi.nlm.nih.gov/geo/query/acc.cgi?acc=GSE153164 |
| Mouse developing visual cortex scRNA-seq | Gao et al.<sup>31</sup> | https://allen-developmental-mouse-atlas.s3.amazonaws.com/scRNA/DevVIS_scRNA_processed.h5ad |
| Ferret developing cortex scRNA-seq | Lucia Del-Valle-Anton et al.<sup>33</sup> | GEO GSE234305,<br>https://www.ncbi.nlm.nih.gov/geo/query/acc.cgi?acc=GSE234305 |
| Ferret developing cortex scRNA-seq | Merve Bilgic et al.<sup>32</sup> | https://ddbj.nig.ac.jp/public/ddbj_databases/dra/fastq/DRA016/DRA016867/ |
| **Experimental models: Cell lines** |  |  |
| Human: H9 hESC line | Thomson et al.<sup>123</sup> | hPSCreg ID: WAe009-A |
| Chimpanzee (*Pan troglodytes*): MARI iPSC line | Kitajima et al.<sup>103</sup> | Courtesy from Masanori Imamura lab, Kyoto University |
| Chimpanzee (*Pan troglodytes*): IKU iPSC line | Lin et al.<sup>120</sup> | Courtesy from Masanori Imamura lab, Kyoto University |
| Gorilla (*Gorilla gorilla*): WER1 iPSC line | Generated by the Masanori Imamura laboratory | Courtesy from Masanori Imamura lab, Kyoto University |
| **Software and algorithms** |  |  |
| Original analysis code for trajectory reconstruction, divergence metrics, branch-resolved contrast, gene clustering, and spatial neighbourhood modelling | This paper | https://github.com/XD-Sheu/Coupling-divergence_2026; Zenodo: https://doi.org/10.5281/zenodo.22026149 |
| Seurat v5.0 | Hao et al.<sup>104</sup> | https://satijalab.org/seurat/; RRID: SCR_007322 |
| dynamo v1.3 | Qiu et al.<sup>105</sup> | https://github.com/aristoteleo/dynamo-release |
| Scanpy v1.12 | Wolf et al.<sup>106</sup> | https://scanpy.readthedocs.io/; RRID: SCR_018139 |
| Imaris v9.0 | Oxford Instruments (Bitplane) | https://imaris.oxinst.com/; RRID: SCR_007370 |
| Fiji/ImageJ v1.54 | Schindelin et al.<sup>107</sup> | https://fiji.sc/; RRID: SCR_002285 |
| WGCNA v1.74 | Langfelder and Horvath<sup>108</sup> | https://cran.r-project.org/package=WGCNA; RRID: SCR_003302 |
| **Other** |  |  |
| TrajMammal, interactive trajectory-resolved gene expression browser | This paper | https://trajmammals.com |

### Method details

#### Neurodevelopmental Datasets, Quality Control and Integration

To characterize the expression dynamics of orthologs (1:1 type) across mammalian cortical development, we curated publicly available single-cell RNA sequencing (scRNA-seq) datasets from the developing somatosensory cortex of human^28^, rhesus macaque^29^, mouse^30,31^ and ferret^32,33^. Developmental stages were selected to encompass comparable periods spanning cortical development. All downstream analyses were conducted using Seurat v5^109,110^ in R. Cells were retained if they satisfied the following quality control criteria: total UMI counts (nCount_RNA) > 1,000, detected genes (nFeature_RNA) > 1,000, log_10_(genes per UMI) > 0.9, and mitochondrial transcript fraction < 25%. Cells failing to meet these thresholds were excluded from further analysis. Normalization and variance stabilization were performed using SCTransform^111^, with regression of sequencing depth, gene complexity, mitochondrial content to mitigate technical and biological confounders unrelated to lineage identity. Following the method described before^29^, highly variable genes (HVGs) were identified per batch via Seurat’s variance-stabilizing transformation; to minimize integration bias across developmental stages, samples were grouped into three transcriptomically comparable temporal windows (early, middle, late). HVGs were selected using SelectIntegrationFeatures across groups, and the union of selected genes was used for downstream dimensionality reduction and integration.

#### Dimensionality Reduction, Clustering, and Cell Type Annotation

Principal component analysis (PCA) was performed on scaled gene expression matrices, and the first 30 principal components were retained based on inspection of the elbow plot and variance explained. Batch correction and dataset integration were performed in reduced-dimensional space using Harmony^112^, fastMNN^113^, canonical correlation analysis (CCA) ^109^, and reciprocal PCA (RPCA) ^109^, allowing comparative evaluation of integration performance. We applied reciprocal PCA for downstream analysis because it preserves inter-species heterogeneity within each cluster in the integrated dataset. The full analytical code including other integration trails is available at https://github.com/XD-Sheu/Coupling-divergence_2026. Uniform Manifold Approximation and Projection (UMAP)^114^ was applied to the RPCA embeddings using default parameters for visualization. Unless otherwise specified, downstream analyses were conducted using expression values from the SCT assay, in which sequencing depth and batch effects had been regressed out. Cell clustering was performed using the Louvain community detection algorithm^115^ on the first 30 integrated dimensions, with k-nearest neighbors set to 25 and a resolution parameter of 1.4. Cluster identities were assigned based on differential gene expression analysis using the Wilcoxon rank-sum test with Bonferroni correction^116^. For each cluster, the top 50 differentially expressed genes were ranked by –log_10_(adjusted p-value) and filtered for >30% within-cluster expression and >0.5 log2 fold change relative to all other cells. Cluster annotation was further refined using established canonical marker genes. Seven primary cell types were identified and defined by the following canonical markers: ventricular radial glia (*HMGA2, HES1, EMX2*), outer radial glia (*HOPX, TNC, LIFR*), truncated radial glia (*CRYAB. FSTL1, ANXA1*), intermediate progenitor cells to immature excitatory neurons (*EOMES, NEUROD2, ELAVL4*), Glia intermediate progenitors (*EGFR, SPRY2, F3*), astrocytes (*GFAP, ALDH1L1, AQP4*), and oligodendrocyte precursor cells to oligodendrocyte (*PDGFRA, OLIG2, SOX10*). Each cell type was subsequently subsetted for downstream species-comparative analyses.

#### Local Topology Construction, Transition Matrix and Velocity Projection

To infer the developmental trajectories of ventricular progenitor cells and their downstream lineages, we first calculated a per-cell pseudotime score using Monocle ^117,118^. A principal graph was then fitted to these UMAP coordinates using reversed graph embedding to model the trajectory backbone. To establish the origin of the trajectory, the root of the pseudotime axis was explicitly defined by selecting the cluster enriched with the youngest ventricular progenitor cells, determined by average developmental stage. The geodesic distance along the principal graph from this root was extracted as the per-cell pseudotime score. This calculation was implemented independently for each species-specific dataset utilized in this analysis. To translate the scalar pseudotime scores into directional cell-wise velocities, we first established the local cellular topology. A -Nearest Neighbors (kNN) graph was constructed in the principal component analysis (PCA) space. This step was strictly topological; it defined the unweighted connectivity and permissible transition edges between transcriptionally adjacent cells, masking out distant cells to restrict velocity calculations to local neighborhoods.

Using the defined kNN topology and the Monocle-derived pseudotime scores, a transition matrix was constructed utilizing principles of discrete differential geometry (Hodge theory) implemented in Dynamo^105^. A discrete gradient operator was applied to the graph’s incidence matrix to calculate the localized flow of pseudotime. Specifically, the scalar difference in pseudotime was computed across every valid edge, effectively defining the developmental gradient without relying on raw Euclidean distance weights. Finally, these edge-specific gradients were compiled into a transition matrix and projected into low-dimensional embeddings to yield cell-wise velocity vectors. For each cell, the transition weights were aggregated with the spatial displacement vectors of its connected neighbors. This projection was performed onto the PCA space, allowing for the multidimensional calculation.

#### Continuous Vector Field Reconstruction and Vector Calculus

To transition from discrete, cell-wise developmental velocities to a global, continuous mathematical model, we reconstructed a differentiable vector field using the Sparse Vector Field Consensus (SparseVFC) algorithm^119^. Given the cell coordinates *X* in the PCA space and their associated discrete velocity vectors *V* (derived from the transition matrix), SparseVFC learns a continuous vector field function *f* such that:

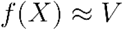

To ensure computational tractability while maintaining the global topology of the dataset, we utilized a sparse approximation leveraging M = 15000 control points. This approach maps the functional space within a Reproducing Kernel Hilbert Space (RKHS) using a Gaussian kernel, significantly reducing computational complexity compared to dense matrix inversions. The algorithm iteratively fits the field, robustly down-weighting outlier vectors and evaluating the cosine correlation between observed and predicted velocities to prevent local distortions.

#### Least Action Path (LAP) and Cell Fate Transition Modeling

To mathematically model the specific transition trajectories between distinct neurodevelopmental cell types (e.g., vRG, oRG and mature states), we first identified representative source and target cells. Reference coordinates corresponding to the phenotypic extremes of each cell type were defined within the UMAP embedding. The exact starting and ending cell indices for the transitions were subsequently assigned by identifying the nearest neighbor cells to these coordinate points. To initialize the optimization algorithm, an initial coarse path was estimated by calculating the shortest path connecting the source and target cells through the kNN transition graph.

We then utilized the continuous vector field function *f*(*x*) learned in the PCA space to optimise the predicted path. We quantified the transition probability using the Onsager-Machlup action, where the action *S* over a path of length *T* is defined as the deviation of the cell’s actual velocity 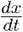 from the deterministic vector field *f*(*x*):

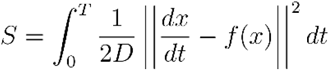

where *D* represents the diffusion coefficient (stochastic noise).In practice, the algorithm discretizes the path into *N* waypoints. The discrete action was minimized using the analytical Jacobian matrix of the vector field:

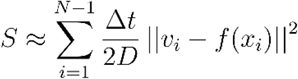

where *x*_*i*_ is the midpoint between consecutive states, *v*_*i*_ is the discrete velocity across the step, and Δ*t* is the optimized time step. We iteratively minimized this action functional utilizing the least_action function in Dynamo, executing Expectation-Maximization (EM) steps to co-optimize the spatial path coordinates and the transition time step Δ*t*, yielding the optimal Least Action Path (LAP) in the PCA space.

#### Divergence metrics on developmental trajectories

Trajectory expression matrices were returned from log space to linear space before any distance was computed, so that observed values and null distributions were built on a common scale. For gene in species *s, xi,s,t,u* let denote expression at step *u* of trajectory *t*, with *t* = 1,…,13 and *u* = 1, …,27. Two nested distributions were derived from this matrix.

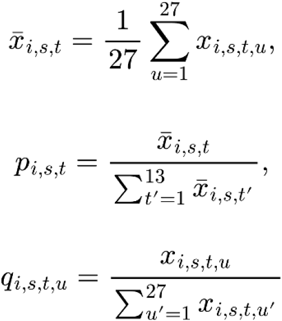

The allocation distribution is the mean expression across the 27 steps of each trajectory, normalised across the 13 trajectories, and describes which lineages a gene is used on independently of its overall expression level. The progression distribution is the 27-step profile within a single trajectory, normalised within that trajectory, and describes where along a lineage a gene is used independently of how much expression that lineage carries.

Both distributions were smoothed with a symmetric Dirichlet pseudocount, 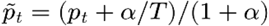, with alpha = 0.05 and ***T*** the number of categories (13 for allocation, 27 for progression). Smoothing suppresses low-count noise and places a strictly positive floor on every category, so all quantities below are defined for every one of the 9,003 one-to-one orthologs.

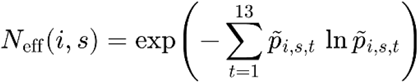

Lineage broadness (***N*** eff, above equation) is the exponential of the Shannon entropy of the smoothed allocation distribution, interpretable as the effective number of trajectories a gene occupies. It equals 1 for a gene confined to a single trajectory and 13 for a gene distributed evenly across all of them, and was averaged over the four species unless stated otherwise.

Allocation divergence between species ***a*** and ***b*** is the Jensen-Shannon distance between their allocation distributions,

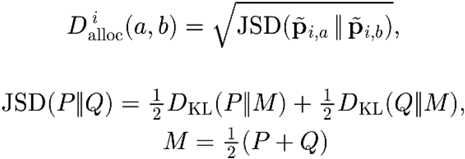

that is the square root of the Jensen-Shannon divergence taken with natural logarithms. It is bounded in [0, 1] and satisfies the triangle inequality, so values are comparable across species pairs.

Progression divergence was computed within each trajectory and then aggregated across trajectories.

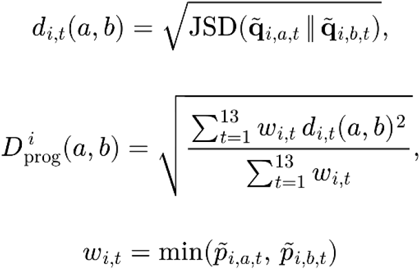

The per-trajectory distance is the Jensen-Shannon distance between the two species’ progression distributions for that trajectory. Distances were combined on the squared scale with weights 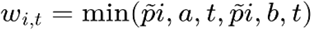, so that each trajectory contributes in proportion to the expression the two species share on it and trajectories on which neither species deploys the gene contribute negligibly. Because the weights derive from smoothed proportions they are strictly positive, and progression divergence was therefore defined for every gene and every species pair.

Two alternative aggregations were computed for robustness. The unweighted variant replaces the weights with a constant, giving the root mean square of the 13 per-trajectory distances. The consensus variant uses

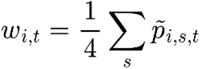

the mean allocation across all four species. The consensus weight does not depend on the species pair being compared, which preserves the triangle inequality. Aggregated progression divergence is accordingly reported as a divergence rather than as a distance, and per-gene divergence trees were built from the consensus variant for this reason. Each metric yields six values per gene, one for each species pair. These were summarised by their maximum in the main analyses and by their mean in the corresponding supplementary analyses.

#### Expression magnitude, detection threshold, and deconfounding

Expression magnitude for a gene in one species was its mean expression across all trajectories and steps, summarised across species by the maximum.

An expression-matched null was used to establish the range over which divergence reflects common ancestry rather than sampling. Genes were assigned to 25 quantile bins of log_10_ expression magnitude, and within each bin gene labels were permuted without fixed points, pairing every gene with a different gene of comparable expression. Both metrics were then recomputed between gene *i* in species ***a*** and its assigned partner in species ***b***, for all six pairs, and summarised by the same statistic used for the observed values. This procedure breaks orthology while holding expression constant, so the difference between observed and null isolates the component of cross-species similarity attributable to shared ancestry. Observed and null values were smoothed against log_10_ magnitude by LOESS (span 0.6). The deviation crossed zero at a magnitude of 0.023, which was taken as the detection threshold, and the 7,212 orthologs above it were retained for all downstream analysis.

To remove expression magnitude analytically, each metric was regressed on log_10_ magnitude using a natural cubic spline with four degrees of freedom and replaced by the residual. Residualisation was applied identically to both axes, so that the correlation between residuals is the partial correlation of the two metrics given expression magnitude. The same relationship was also examined within deciles of expression magnitude, computing the correlation separately in each decile with 500 gene-bootstrap replicates per decile for confidence intervals.

The dependence of coupling on divergence time was assessed within species pairs. For each pair, the Pearson correlation was computed between that pair’s own allocation and progression divergence. Correlations were transformed with Fisher’s *z*, with confidence intervals given by 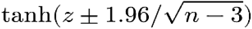, and regressed on divergence time by weighted least squares with weights *n* − 3. Divergence times were taken as 29 My for human and macaque, 87 My for the two primate to mouse pairs, and 94 My for the three pairs involving ferret.

#### Conserved lineage-specifying genes

Genes falling below the 10th percentile on allocation divergence, progression divergence, and ***N*** eff simultaneously were retained as conserved lineage-specifying genes (n = 87). Each gene was assigned to the trajectory carrying the largest share of its consensus allocation, defined as the mean allocation distribution across the four species (**Figure 2F**). Assignments were curated against the literature into three categories: fate determinants, defined as genes whose misexpression is sufficient to induce the corresponding identity; lineage-associated genes, with documented expression or marker status on that lineage but no reported instructive role; and genes with no reported role on the assigned lineage (**Supplementary Table S2**).

#### Branch-specific contrasts and species-biased gene sets

To assign divergence to a single branch, the two primates were treated as an ingroup and the two remaining species as outgroups. For each gene the ingroup distance is the divergence between human and macaque, and the contrast is the mean human-to-outgroup distance minus the mean macaque-to-outgroup distance.

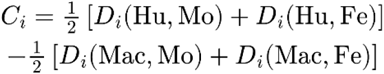

A positive contrast places divergence on the human branch subsequent to the human and macaque split, and the two diagonals in **Figures 3D** and **3E** bound the region in which divergence is attributable entirely to one branch. The contrast was computed separately on allocation divergence and on progression divergence, and the genes with the largest positive values on each (n = 150) were taken as human-biased reallocation genes and human-biased retiming genes respectively. The same construction was applied with each species in turn as the focal branch, using its nearest relative among the sampled species as the comparison branch and the remaining two species as outgroups (measured species-biased contrast are in **Supplementary Table S1**).

Enrichment of the overlap between the two gene sets was assessed by permutation. Gene labels on the progression contrast were shuffled, the top 150 genes were re-selected, and the size of the overlap with the unshuffled top 150 reallocation genes was recorded; this was repeated 1,000 times for each focal species, and the mean of the resulting null distribution is reported. Shuffling preserves both marginal contrast distributions and removes only the correspondence between them, so the comparison tests co-membership rather than the shape of either ranking.

Per-gene divergence trees were constructed from the six pairwise divergences for a single gene, assembled into a four by four distance matrix and fitted by BIONJ. Trees were built separately from allocation divergence and from consensus-weighted progression divergence. A per-gene tree carries no gene-level bootstrap support, and node support is therefore not reported. Any negative branch lengths arising from non-additive distances were set to zero.

#### Cortical organoid cultivation and immunofluorescence

Cerebral organoids were generated from human embryonic stem cells (line #H9 or WA09^123^), chimpanzee iPSCs (lines #mari and #iku^120^) and gorilla iPSCs (lines #Wer1) using the STEMdiff™ Cerebral Organoid Kit (STEMCELL Technologies, #08570) according to the manufacturer’s instructions, with minor modifications based on method described before^121,122^. Briefly, after removing differentiated regions from pluripotent stem cells cultures, cells were dissociated with accutase enzyme solution (Sigma-Aldrich, catalog #A6964), resuspended in EB formation medium supplemented with 10 µM Y-27632 (STEMCELL Technologies, Catalog #72302), and seeded at 5000 cells/well in 96-well ultra-low attachment plates (Corning, Catalog #7007). Plates were then incubated at 37°C, with EB formation medium replenished on every other day. On day 5, EBs were transferred to 24-well plates (coated with Anti-Adherence Rinsing Solution, Catalog #07010) in an induction medium and incubated for 48 h to form neuroepithelium. On day 7, EBs were embedded in Matrigel droplets (BD, Cat# 354277), polymerized, and transferred to 6-well plates (coated with Anti-Adherence Rinsing Solution) in Expansion Medium for 3 days to develop budding neuroepithelia. On day 10, medium was replaced with maturation medium, and organoids were cultured on an orbital shaker at 37°C with medium changes every 2 days. e intensity thresholding; local contrast region-growing; background subtraction disabled).

At two months of culture (DIV60) and late stages (DIV160), cortical organoids were washed three times for 10 minutes each with D-PBS and fixed overnight in freshly prepared 4% paraformaldehyde (PFA) at 2–8°C. Organoids were cryoprotected in 30% sucrose in D-PBS overnight at 4°C until equilibrated, infiltrated with gelatin solution (7.5% gelatin and 10% sucrose in D-PBS) at 37°C for 1 h, transferred to embedding molds, snap-frozen in a dry ice/ethanol slurry, and stored at -80°C. Frozen blocks were equilibrated in a cryostat for 30 min and sectioned at 15 µm thickness onto slides. Sections were air-dried at room temperature, washed with PBS-T at 37°C for 10 min and blocked for 1 h at room temperature in a humidified chamber with blocking solution (5% normal donkey serum in PBS-T). Sections were then incubated overnight at room temperature in a humidified chamber with primary antibodies diluted in primary dilution buffer (PBS-T containing 0.05% sodium azide and 5% BSA): goat anti-PDGFRα (R&D Systems AF1062, 1:500), mouse anti-SOX10 (Abcam ab216020, clone SOX10/1074; 1:500), rabbit anti-PSD-95 (Proteintech 20665-1-AP, 1:300), and rabbit anti-GluN2B/GRIN2B (Proteintech 21920-1-AP, 1:300). After three PBS-T washes, sections were incubated for 2 h at room temperature in a humidified chamber with secondary antibodies diluted in PBS-T: Alexa Fluor 488 donkey anti-rabbit IgG (H+L) (1:1000; Abcam, ab150073), Alexa Fluor 555 donkey anti-mouse IgG (H+L) (1:1000; Abcam, ab150106), Alexa Fluor 647 donkey anti-goat IgG (H+L) (1:1000; Abcam, ab150131). Following three additional 30-min PBS-T washes, slides were air-dried for 5 min, mounted on glass slides using DAKO fluorescent mounting medium. Imaging was performed using ZEISS LSM 900 and LSM 990/Airyscan2 Multiplex confocal microscopes. Z-stacked images were acquired using 60x and 100x magnification objectives with a 0.3-0.5 µm z-interval for subsequent 3D reconstruction.

#### Volumetric Reconstruction and Puncta Quantification

Volumetric reconstructions and quantitative analyses of the fluorescence z-stacks were performed using Imaris software (Bitplane). To define the cellular volume of OPCs, the PDGFRα fluorescence channel was reconstructed using the “Surface” module (surface grain size: 0.25 µm; absolute intensity thresholding; local contrast region-growing; background subtraction disabled).

Synaptic proteins were reconstructed using the Imaris “Spots” module, initialized with an estimated diameter of 0.2 µm and background subtraction enabled. The local contrast region-growing algorithm was applied to accurately capture the volume of each synaptic punctum. The shortest distance from the center of each spot to the generated PDGFRα cell surface was calculated. GluN2B puncta located within 0.5 µm of the external cell surface, as well as all internalized puncta, were retained. PSD-95 puncta were strictly filtered to include only internalized puncta located inside the PDGFRα surface. Finally, the expression of each synaptic protein was quantified using two normalized metrics: (1) the synaptic signal volume fraction, defined as the total volume of the filtered spots divided by the total PDGFRα-positive cell volume; and (2) the puncta density, calculated as the absolute count of filtered spots per unit of cell volume.

#### Temporal Clustering of Synaptic Genes

To analyze the broader synaptic program, a 601-gene panel was compiled from synaptic-transmission Gene Ontology terms (GO:0050804, GO:0015459, GO:0017080, GO:0051932). Mean expression trajectories were compared between *in vivo* human (GW14 to adulthood)^28^ and mouse (E15 to P56)^31^ OPCs. To resolve specific temporal patterns, the 601 genes were clustered based on their joint human and mouse trajectories into four temporal modules (M1–M4) using the WGCNA package in *R*. Network construction was performed using the blockwiseModules function with the following parameters optimized for distinct temporal dynamics: power = 6, TOMType = “signed Nowick 2”, networkType = “signed”, minModuleSize = 30, and mergeCutHeight = 0.25.

#### Single-Cell Module Scoring and Differential Expression

Transcriptional heterogeneity among early human *in vivo* OPCs (GW14–GW18; n = 231 cells) was evaluated by calculating a Cluster 2 expression score for each cell using the AddModuleScore function in Seurat. The population was dichotomized into “high-score” (top 22% of the distribution; module score > 0.465) and “low-score” OPCs. After confirming that baseline core OPC markers (PDGFRA, SOX10) were equivalently expressed between the two groups, differential gene expression and subsequent GO enrichment analyses were performed to characterize the distinct functional states (e.g., neuronal vs. proliferative) separating the subpopulations.

#### Re-analysis of the public MERFISH atlas

We obtained the MERFISH dataset of the developing human cortex from Qian et al.^60^ and Wang et al.^28^, comprising 16.3 million cells across 8 developmental stages (GW15: 4.7 M cells from samples *UMB1117* and *UMB1367*; GW20: 7.7 M from *FB080* and *FB121*; GW22: 2.3 M from *FB123*; GW24: 0.2 M from *ARKFrozen-62-PFC, ARKFrozen-65-V1*; GW33: 0.05 M from *NIH-5900-BA17*; GW34: 1.1 M from *UMB5900*, GW35: 0.05 M from *NIH-4365-BA10;* Infancy 8 month: 0.02M from *NIH-4392-BA17*). The authors’ three-level hierarchical cell-type annotations (*H1, H2, H3*, in Qian et al.) as well as “type_updated” in Wang et al. were retained without further re-clustering. Counts were depth-normalized to 10,000 per cell with scanpy.pp.normalize_total and log1p-transformed with scanpy.pp.log1p prior to all downstream analyses. For the focal-receiver analyses we re-aggregated the original annotations into two combined populations: an oRG-group comprising cells with *H2* = *oRG1* together with *H3* ∈ {*oRG-1, oRG-2, oRG-3, oRG-4*}, and an EN-oSVZ-group comprising *H2* ∈ {*EN-oSVZ-1, En-oSVZ-2*}. For the multi-sender regression below we additionally constructed a composite sender label, sender_composite, which used the OPC subtype value (*neuronal-OPC, proliferative-OPC*) for cells with *H2* = *OPC*, and the *H2* annotation for all other cells.

#### Spatial neighbor graph and Neighborhood composition

Per imaged section (per *sample × region* combination), we constructed a Delaunay triangulation of cell centroids using squidpy.gr.spatial_neighbors, yielding a binary adjacency matrix *A* ∈ 0,1^*N*×*N*^ in *A*_*ij*_ = 1 which if cells ***i*** and ***j*** share a Delaunay edge.

For each *i* cell and each cell-type label *t*, we computed the fraction of *i*’s graph neighbors of type *t*.

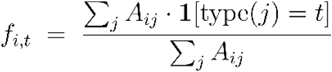

Stage-stratified neighborhood enrichment for OPCs (**Figure 6**) was computed by averaging *f*_*i,t*_ over all OPC cells *i* within a developmental stage and comparing to the cohort-wide mean abundance of type *t*.

#### Multi-sender ordinary least squares regression

For each receiver cell type involved in oRG and EN-oSVZ category, restricted to GW15 cells, and each gene *g* expressed in at least 2% of receiver cells, we fit a single multi-sender model,

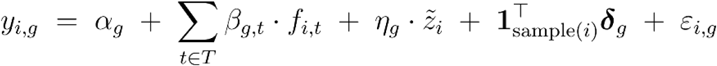

where *y*_*i,g*_ is the log1p-normalized expression of gene *g* in receiver cell *i*; *f*_*i,t*_ is the spatial-neighborhood fraction of sender type *t*, computed from the Delaunay graph using the *sender_composite* label; the sender set *T* comprises *neuronal-OPC, proliferative-OPC, OPC-other*, the union of the top five most-abundant non-OPC cell types in each receiver’s neighborhoods, and a residual “sender-other” pooled bucket; 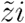 *is z-scored cortical depth;* 1sample(*i*) is the sample-level fixed-effect indicator vector; and *ε*_*i,g*_ is residual error. Models were fit by ordinary least squares with HC3 heteroscedasticity-robust standard errors (statsmodels.api.OLS with cov_type=‘HC3’). Receiver cells with missing 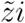 were excluded; per-receiver effective sample sizes are reported in **Supplementary Table S8**. Per-gene p-values were Benjamini– Hochberg FDR-adjusted within each (*R,t*) pair separately. The partial coefficient *β*_*g,t*_ is interpretable as the marginal change in receiver gene-*g* expression per unit increase in the local fraction of sender type *t*, holding all other senders constant.

#### β-distribution comparison and gene-wise classification

For each receiver, we compared the across-gene distributions of *β_{g, neuronal-OPC}* and *β_{g, proliferative-OPC}* using the 1-Wasserstein distance

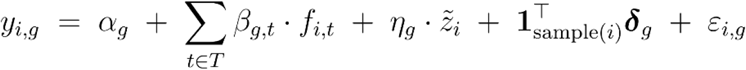

and the two-sample Kolmogorov–Smirnov test, and summarized total coupling magnitude using the L1 and L2 norms

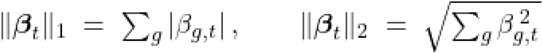

of each per-condition β-vector. For per-gene classification (**Figure 6**), we labeled a gene as neuronal-OPC stronger if *q_{neuronal-OPC}* < 0.05 and |*β_{g, neuronal-OPC}*| *>* |*β_{g, proliferative-OPC}*|, proliferative-OPC stronger if the converse held with *q_{proliferative-OPC}* < 0.05.

#### Permutation test for neuronal-OPC–specific effects

To test whether the per-gene difference Δ*β*_*g*_ = *β*_g,syn-high_ − *β*_g,syn-low_ could arise from random labeling within the OPC pool rather than reflecting genuine OPC-state specific regulation, we permuted the neuronal-OPC and proliferative-OPC labels among OPC cells, stratified by sample (so per-sample counts of each label were preserved), recomputed the neuronal-OPC and proliferative-OPC neighborhood fractions, and refit the multi-sender model. The test was applied to the top 50 candidate genes per receiver, ranked by absoulte value of coefficient value. Across ***n*** perm = 2000, permutations we obtained a per-gene null distribution of 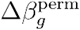. We computed two complementary test statistics: an empirical permutation p-value,

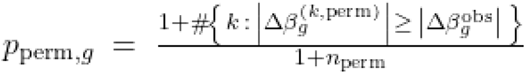

and a parametric Gaussian-approximation p-value

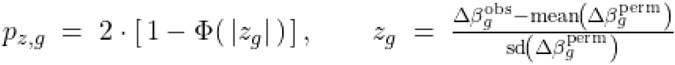

based on the normalized z-score 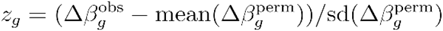. Both p-values were Benjamini– Hochberg FDR-adjusted across the candidate genes; we report the more conservative of the two as *q*_perm_ in the main text. Genes with *q*_perm_ < 0.05 were classified as exhibiting genuine neuronal-OPC–specific coupling.

#### Per-cell ligand–receptor scoring

From a curated panel of 21 ligand–receptor pairs covering glutamatergic (e.g., *SLC17A6*– *GRIK4*/*GRM7*), GABAergic (*GAD2*–*GABRA5*/*GABRB2*/*GABRG3*), synaptic-adhesion (*CLSTN2, IGSF11, LRP4, NECTIN3, SLITRK5*), cadherin (*CDH13*), ECM–integrin (*COL19A1, COL24A1, COL25A1, COL4A5*/*6, COL6A1, SPOCK1* → *ITGA2*), integrin–integrin (*ITGB5*–*ITGA2*), Reelin (*RELN*–*LRP4*), and angiopoietin (*ANGPT2*–*ANGPTL1*) classes, we computed for each focal cell *i* of focal type *F* and each L– R pair (*ℓ, r*) the partner-to-focal contact score.

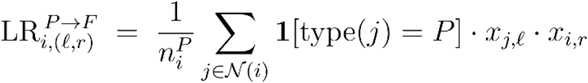

The focal-to-partner score is defined symmetrically. Mean L–R scores per focal subgroup (neuronal-OPC vs proliferative-OPC) were compared using the two-sided Mann–Whitney U test, with Benjamini– Hochberg FDR correction applied across L–R pairs separately within each (partner type, direction) combination.

## Supplementary Figures

**Supplementary Figure S1.**
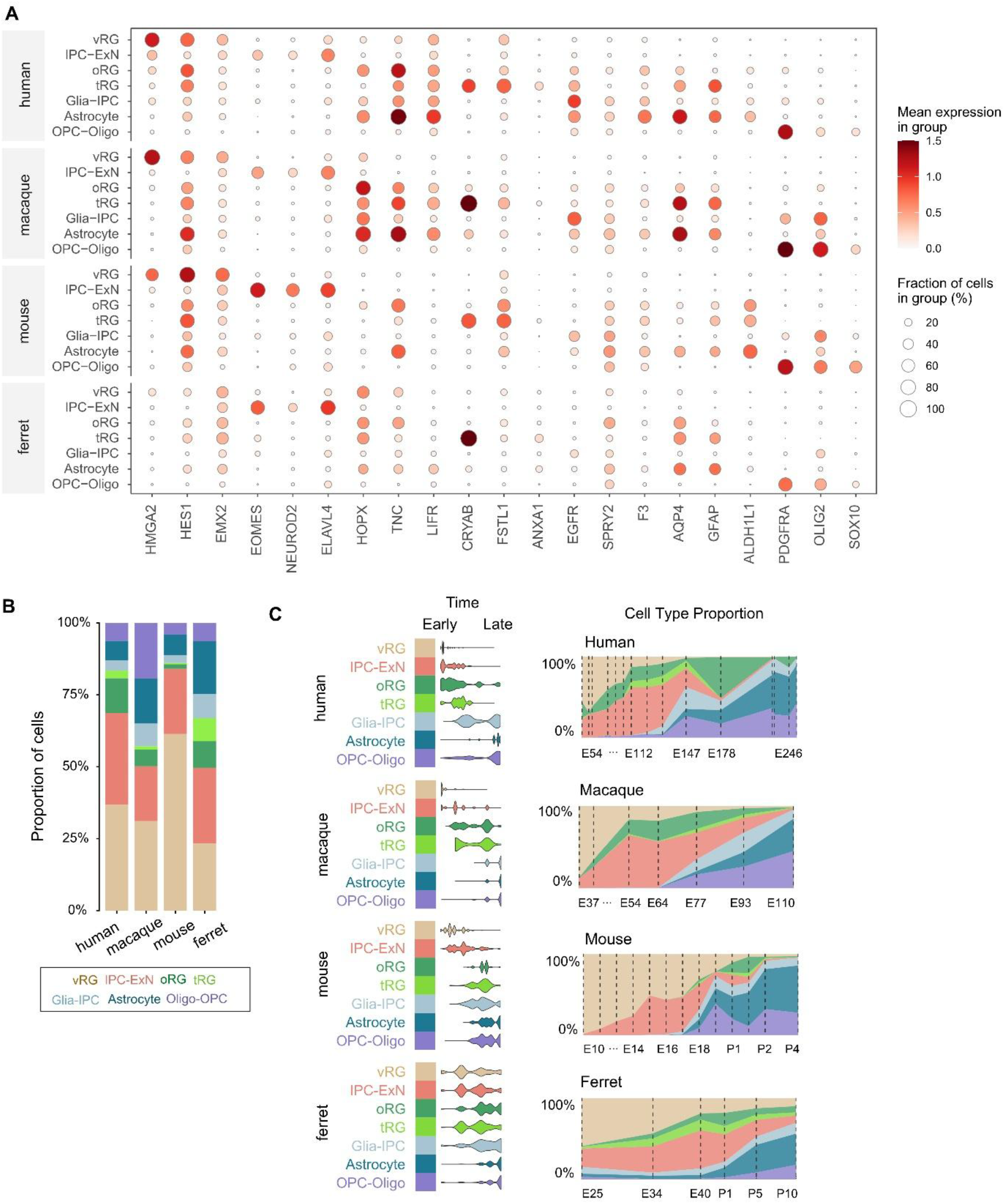
Marker validation and developmental composition of the seven consensus cell types. **A**. Expression of canonical markers across the seven consensus cell types in each species. Dot colour is mean expression within a cell type and dot size the fraction of cells expressing the gene. Markers are ordered by the population they define: *HMGA2, HES1* and *EMX2* for vRG; *EOMES, NEUROD2* and *ELAVL4* for IPC-ExN; *HOPX, TNC* and *LIFR* for oRG; *CRYAB, FSTL1* and *ANXA1* for tRG; *EGFR, SPRY2* and *F3* for glia-IPC; *AQP4, GFAP* and *ALDH1L1* for astrocytes; and *PDGFRA, OLIG2* and *SOX10* for OPC-oligo. **B**. Proportion of cells assigned to each consensus type in each species, across the full sampled window. **C**. Developmental distribution of each cell type. Violin plots (left) show the stages at which each population is sampled, ordered from early to late; stacked area plots (right) show cell-type proportions at each sampled stage, with dashed lines marking the stages.

**Supplementary Figure S2.**
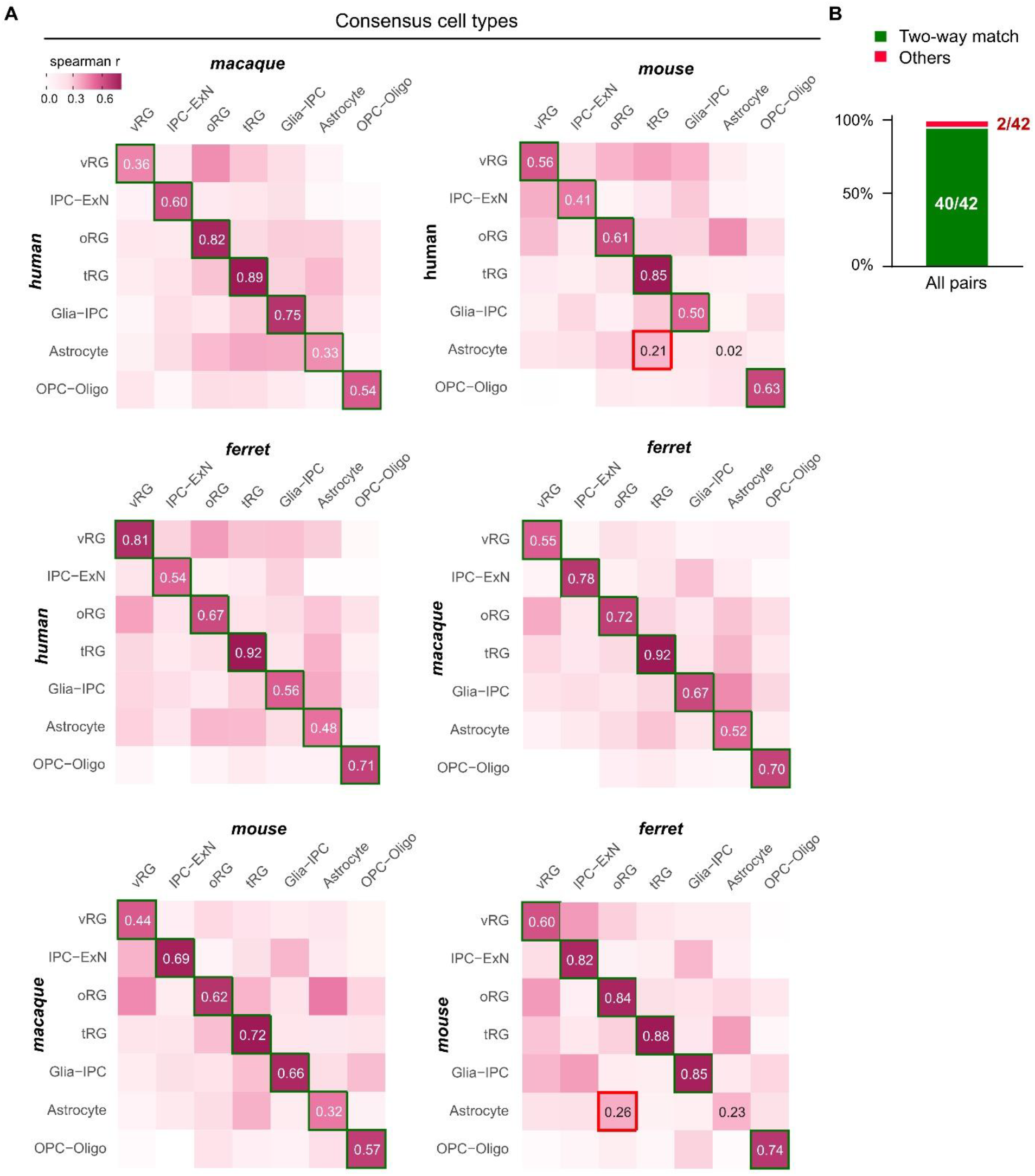
Cell-type correspondence is reciprocal across all six species pairs. **A**. Spearman correlation between the seven consensus cell types in each pair of species, computed on mean expression across one-to-one orthologs (Methods). Green outlines mark reciprocal best matches, in which each type is the highest-correlating partner of the other. Red outlines mark the two comparisons in which the reciprocal criterion was not met, both involving astrocytes: in human against mouse the human astrocyte correlates most strongly with mouse tRG (*r* = 0.21) rather than with mouse astrocyte (*r* = 0.02), and in mouse against ferret the mouse astrocyte correlates most strongly with ferret oRG (*r* = 0.26) rather than with ferret astrocyte (*r* = 0.23). Correlations along the diagonal range from 0.32 to 0.92. **B**. Proportion of reciprocal best matches across all comparisons (40 of 42, 95.2%).

**Supplementary Figure S3.**
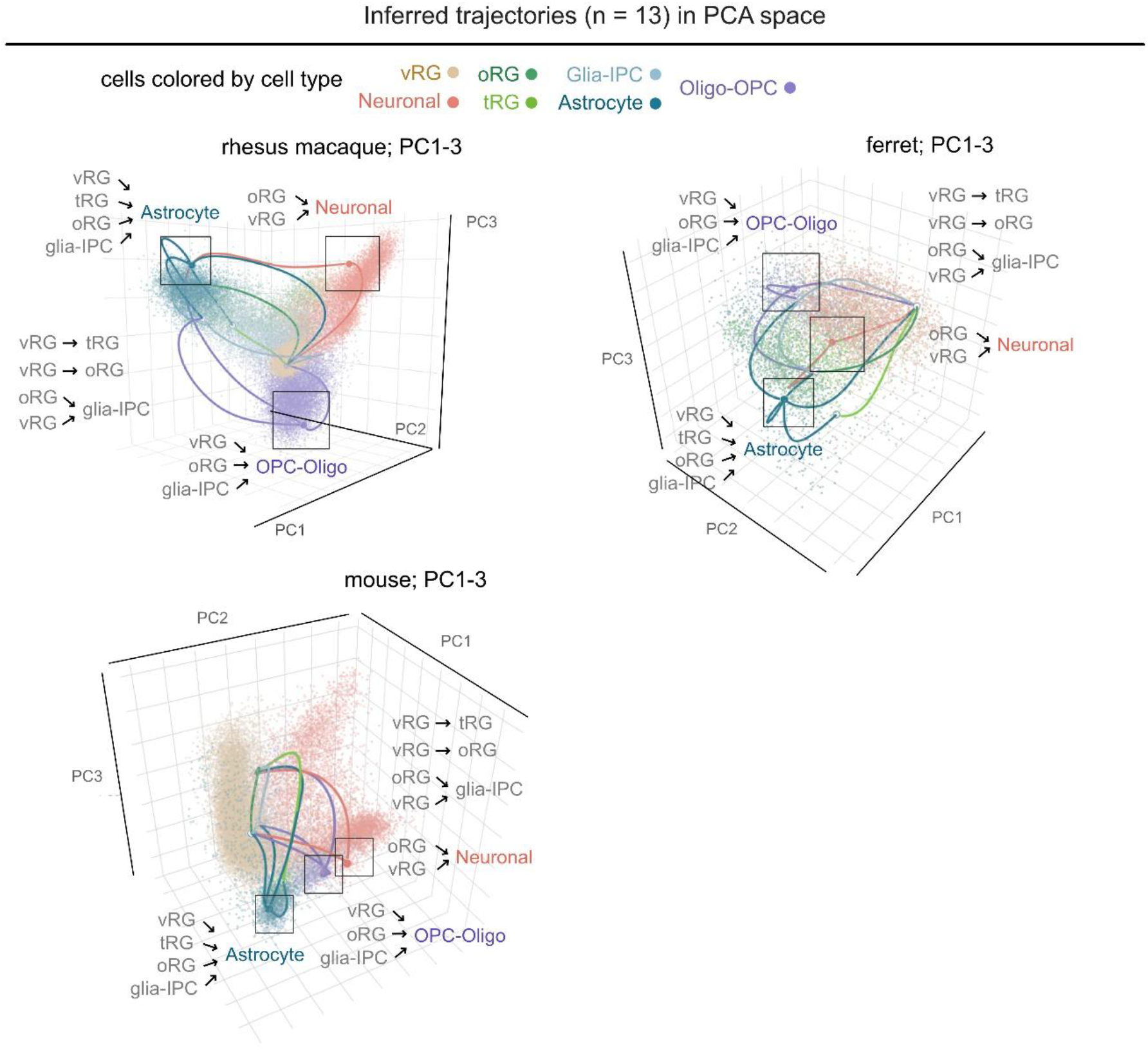
The single-cell developmental trajectories inferred in every species. The 13 inferred trajectories for rhesus macaque, ferret and mouse, projected onto the first three principal components of each species and coloured by consensus cell type. Boxes mark the terminal cell states of the three principal lineage axes, toward neurons, astrocytes and OPC-oligodendrocytes, with the source populations of each trajectory listed alongside. The equivalent projection for human is shown in **Figure 1F**.

**Supplementary Figure S4.**
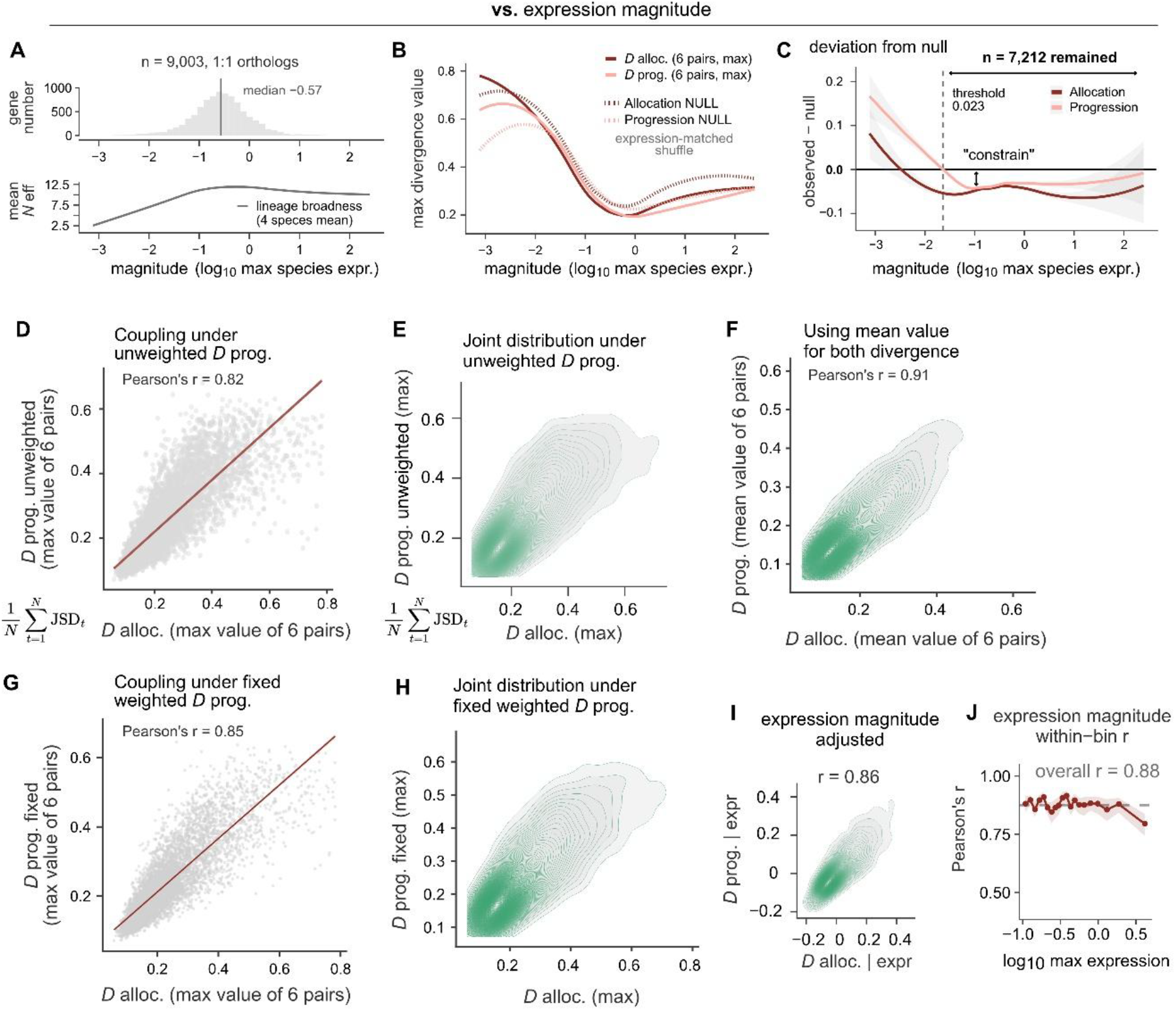
Validation of divergence coupling across expression magnitudes and mathematical controls. **A**. Expression magnitude across the 9,003 one-to-one orthologs (top) and mean lineage broadness against magnitude (bottom). **B**. Allocation and progression divergence against expression magnitude (solid), with expression-matched broken-orthology nulls (dashed). Divergences are summarised as the maximum across the six species pairs. **C**. Observed minus null for both metrics. Values below zero indicate conservation beyond chance. The zero crossing at a maximum species expression of 0.023 defines the detection limit; the 7,212 orthologs above it are retained for all subsequent analyses. **D**. Mathematical control by replacing the mass-weighted aggregation of progression divergence with an unweighted mean across trajectories. **E**. Joint density of the unweighted *D* prog. metrics. **F**. Joint density for *D* alloc. and *D* prog., both calculated from taking mean value (instead of max value) from 6 species-pairs. **G**. Allocation against progression divergence when progression is aggregated with a fixed, pair-independent weighting. *r* = 0.85 (*p* < 2.2 × 10^−16^, n = 7,212). Each trajectory is weighted by the mean allocation across all four species rather than by the mass shared between the pair being compared, which removes the dependence of the weights on the species pair (Methods). **H**. Joint density of the fix weightened *D* prog.. **I**. *D* alloc. and *D* prog. correlation after controlling for expression magnitude for each ortholog. **J**. Pearson correlation between the two metrics computed separately within bins of expression magnitude. Points are per-bin correlations with bootstrap confidence bands; the dashed line marks the correlation across all genes.

**Supplementary Figure S5.**
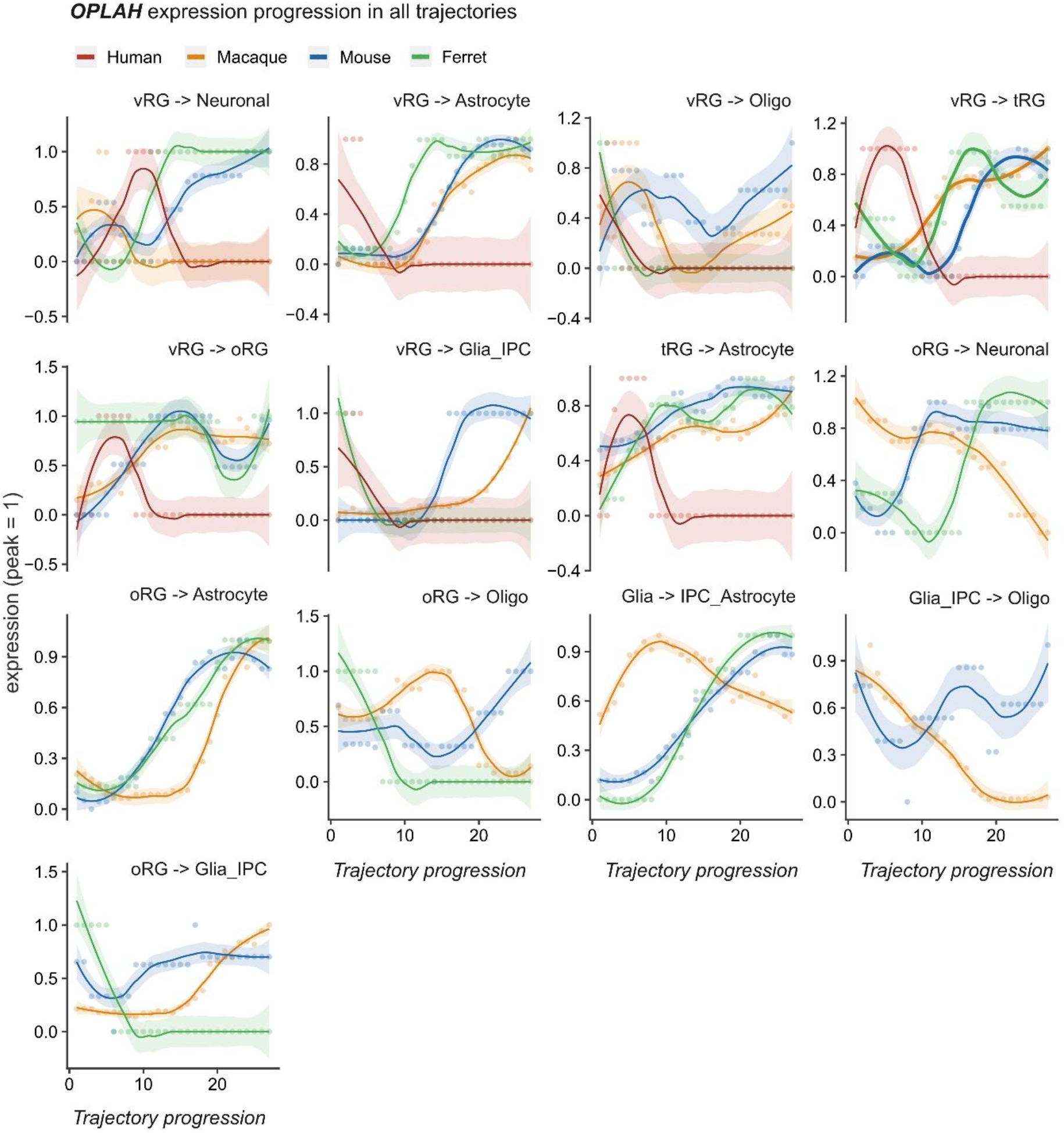
*OPLAH* progression across all trajectories in four species. *OPLAH* expression along each of the 13 inferred trajectories in human, macaque, mouse and ferret. Points are per-step values and lines are LOESS fits with 95% confidence bands, with expression normalised to the peak within each species and trajectory. The tRG to astrocyte trajectory, which carries most of *OPLAH* expression in all four species, is shown in **Figure 3L**. Human expression is confined to the vRG-rooted trajectories and to tRG to astrocyte, and falls to zero on the oRG-rooted and glia-IPC-rooted routes where the other three species retain expression, which is the reallocation quantified in **Figure 3J**. Where human expression is present, it peaks within the first half of the trajectory and declines thereafter, whereas the other species rise across the second half. Normalisation is within species and trajectory, so curve heights are comparable in shape but not in magnitude between panels. Per-step values for all genes and trajectories are available through TrajMammal (**trajmammals.com**).

**Supplementary Figure S6.**
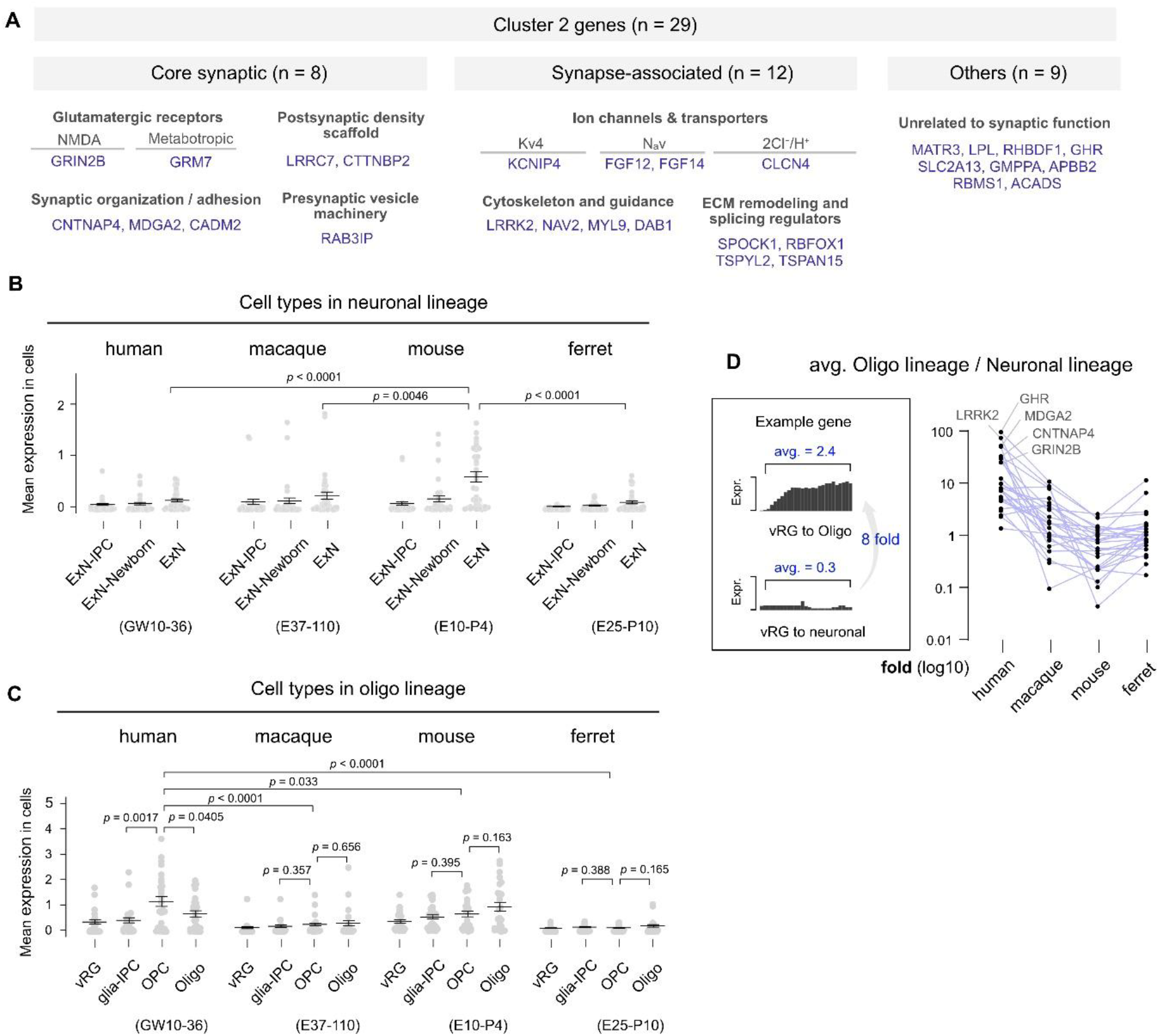
Cluster 2 composition and its expression across the neuronal and oligodendrocyte lineages. **A**. Composition of cluster 2 (n = 29 genes). Twenty of the twenty-nine have documented synaptic roles, comprising eight core synaptic genes across glutamatergic receptors, postsynaptic density scaffolds, synaptic adhesion and presynaptic vesicle machinery, and twelve synapse-associated genes across ion channels, cytoskeletal and guidance regulators, and extracellular matrix and splicing regulators. The remaining nine have no established synaptic role. **B**. Cluster 2 expression across annotated cell types of the neuronal lineage in each species. Each point is the mean expression of one cluster 2 gene within that cell type. Expression is comparable across species over the sampled window, with mouse excitatory neurons above the other three species. **C**. The same analysis across the oligodendrocyte lineage. In human, expression peaks at the OPC stage and declines in mature oligodendrocytes, whereas macaque, mouse and ferret show no significant difference between these stages. Human OPCs exceed the OPCs of every other species. **D**. Ratio of mean cluster 2 expression along the vRG to oligodendrocyte trajectory over the vRG to neuronal trajectory, for each gene in each species. Schematic at left illustrates the calculation for one gene. Every gene is biased toward the oligodendrocyte lineage in human, with the strongest labelled; the bias declines through macaque and mouse and rebounds slightly in ferret. Data in **B** and **C** are mean ± SEM across the 29 cluster 2 genes, with individual genes shown; developmental windows and expression values are those used in **Figures 1 to 3**. Comparisons are Welch’s t-tests with exact *p* values shown on each panel.

**Supplementary Figure S7.**
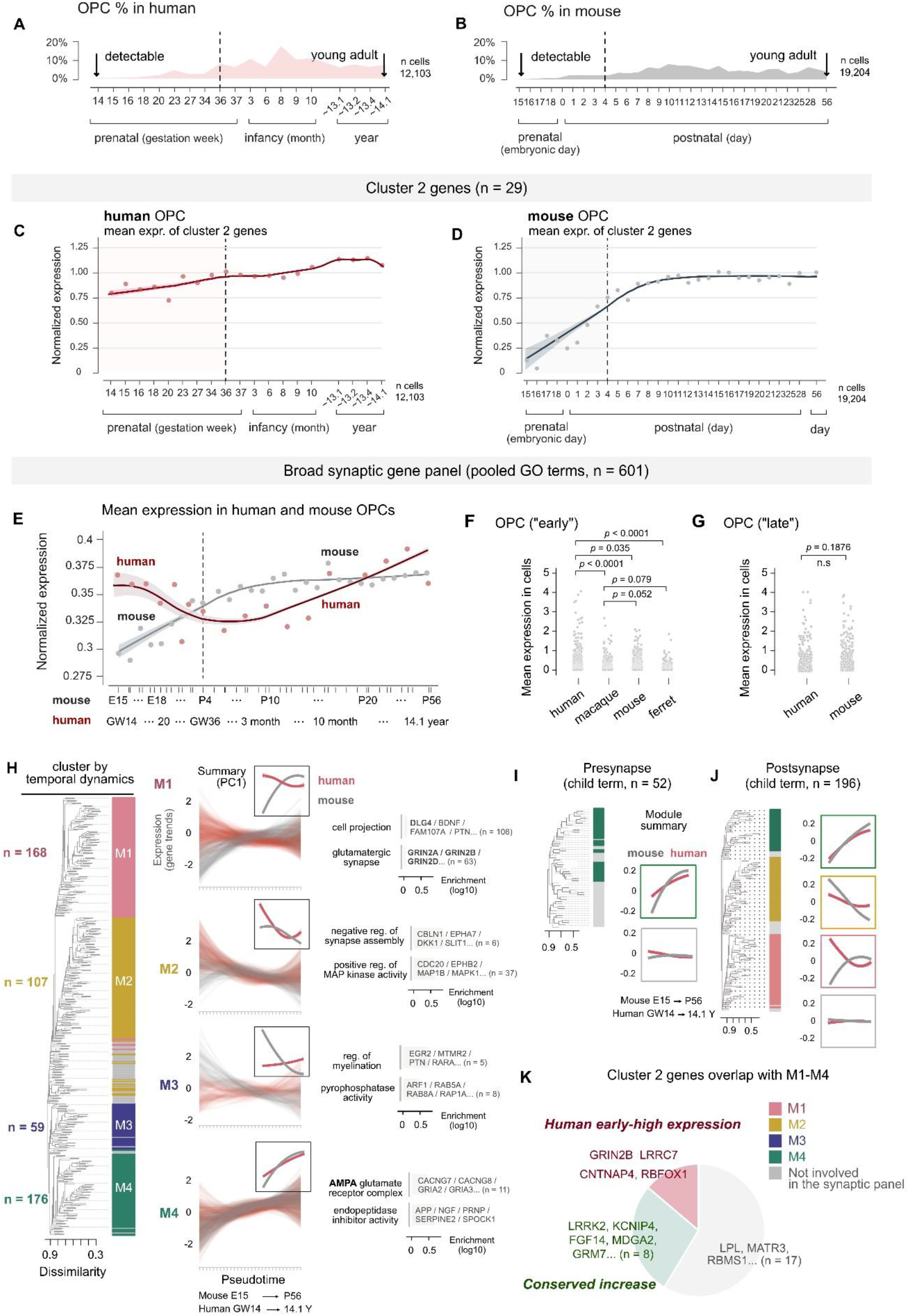
Chronological expression of the synaptic program in human and mouse OPCs. **A-B**. We followed expression within OPCs across chronological development, which is a different axis from the position along a differentiation trajectory analyzed in **Figures 1** to **4**. Human data span GW14 to 14.1 years and mouse E15 to P56. Dashed lines in **A** to **E** mark the boundary between the early window, which corresponds to the developmental range analyzed in **Figures 1** to **4**, and the late window, which is included here only to extend the comparison into postnatal life. Proportion of OPCs among all sampled cells at each stage, in human (**A**) and mouse (**B**). Arrows mark the earliest stage at which OPCs are detectable and the latest stage sampled. **C-D**. Mean cluster 2 expression in OPCs across chronological development in human (**C**) and mouse (**D**). Points are per-stage means and lines are LOESS fits with 95% confidence bands, normalised within species. **E**. Mean expression of a broad synaptic panel of 601 genes, compiled from the synaptic transmission Gene Ontology terms GO:0050804, GO:0015459, GO:0017080 and GO:0051932, in human and mouse OPCs on a common axis. Human expression is highest at the earliest stages and declines across the prenatal window before rising again, whereas mouse expression rises monotonically and reaches the human level only after birth. **F, G**. Panel expression within the early window across all four species (**F**) and within the late window in human and mouse (**G**). Each point is the mean expression of one panel gene. **H**. Clustering of the 601 panel genes by their joint human and mouse temporal dynamics. Dendrogram with module assignments at left; per-gene trends and a first principal component summary for each module in the centre; selected gene ontology enrichments at right. Four modules emerge: M1 (n = 168), in which human declines while mouse rises; M2 (n = 107), declining in human and, after an early rise, in mouse; M3 (n = 59), declining in mouse while human remains flat; and M4 (n = 176), rising in both species. *GRIN2B* and *DLG4* both fall in M1. **I, J**. The same clustering applied separately to the presynaptic (n = 52) and postsynaptic (n = 196) child terms of the panel. A module with the M1 pattern, human high early and declining while mouse rises, is recovered among postsynaptic genes but not among presynaptic genes under identical clustering parameters. **K**. Overlap between cluster 2 and the four temporal modules. Four cluster 2 genes fall in M1, including *GRIN2B*, eight in M4, and the remaining seventeen lie outside the synaptic panel. Data in **F** and **G** are mean ± SEM across panel genes, with individual genes shown; comparisons are Welch’s t-tests with exact *p* values on each panel. Cell numbers are given in **A** and **B**. Panel gene lists and module assignments are in **Supplementary Table S6**.

**Supplementary Figure S8.**
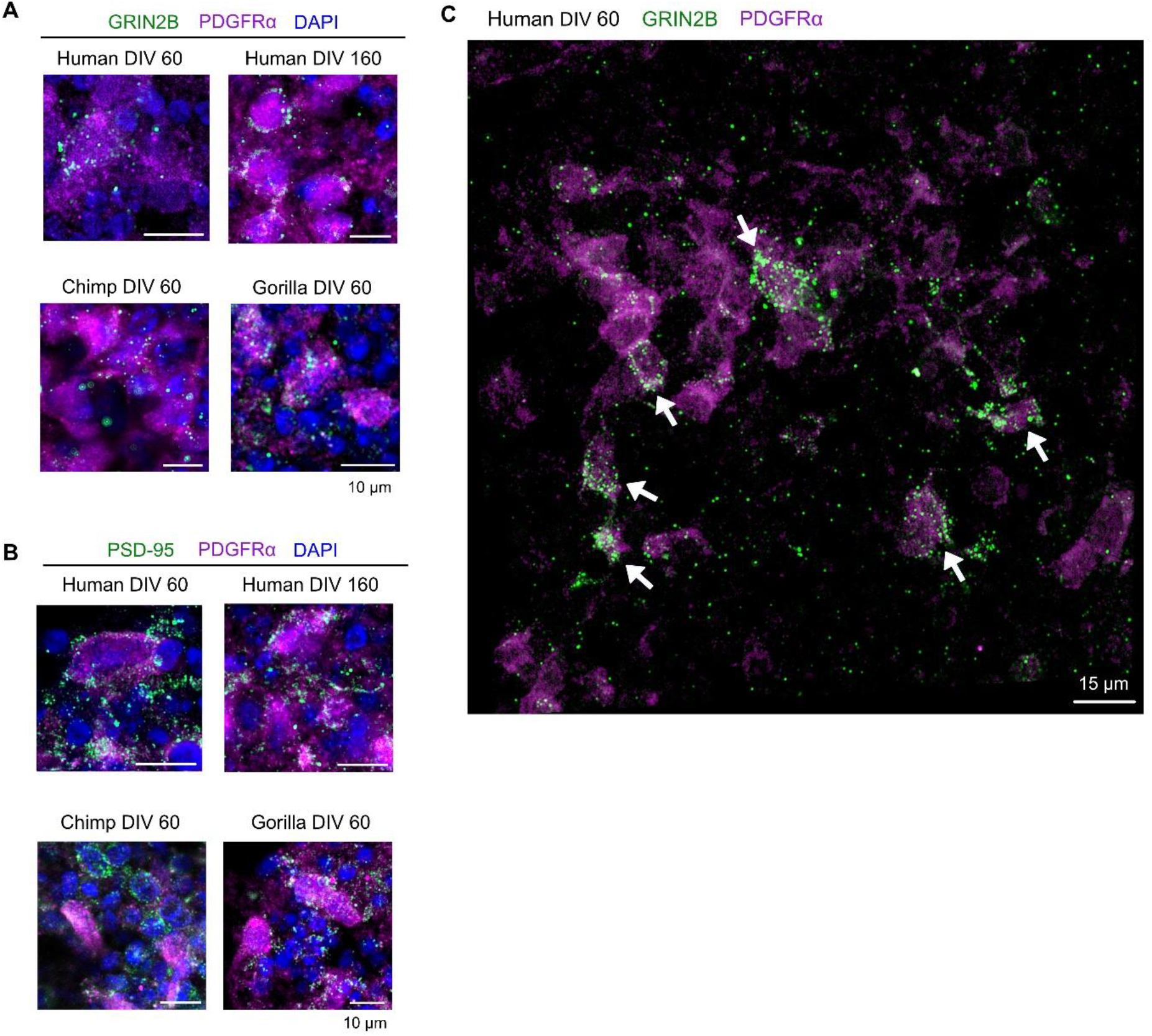
Additional immunofluorescence images of GluN2B and PSD-95 in primate cortical organoids. **A**. Additional fields showing GluN2B (green), PDGFRα (magenta) and DAPI (blue) in human organoids at DIV60 and DIV160 and in chimpanzee and gorilla organoids at DIV60. Quantification of these conditions is given in **Figure 5F** and **5G. B**. The same conditions stained for PSD-95, quantified in **Figure 5I** and **5J. C**. Wide-field view of human organoid at DIV60 stained for GluN2B and PDGFRα. Arrows mark PDGFRα-positive cells carrying dense GluN2B signal, adjacent to PDGFRα-positive cells with signal at or below the level seen in chimpanzee and gorilla. Scale bars are 10 µm in **A** and **B** and 15 µm in **C**.

**Supplementary Figure S9.**
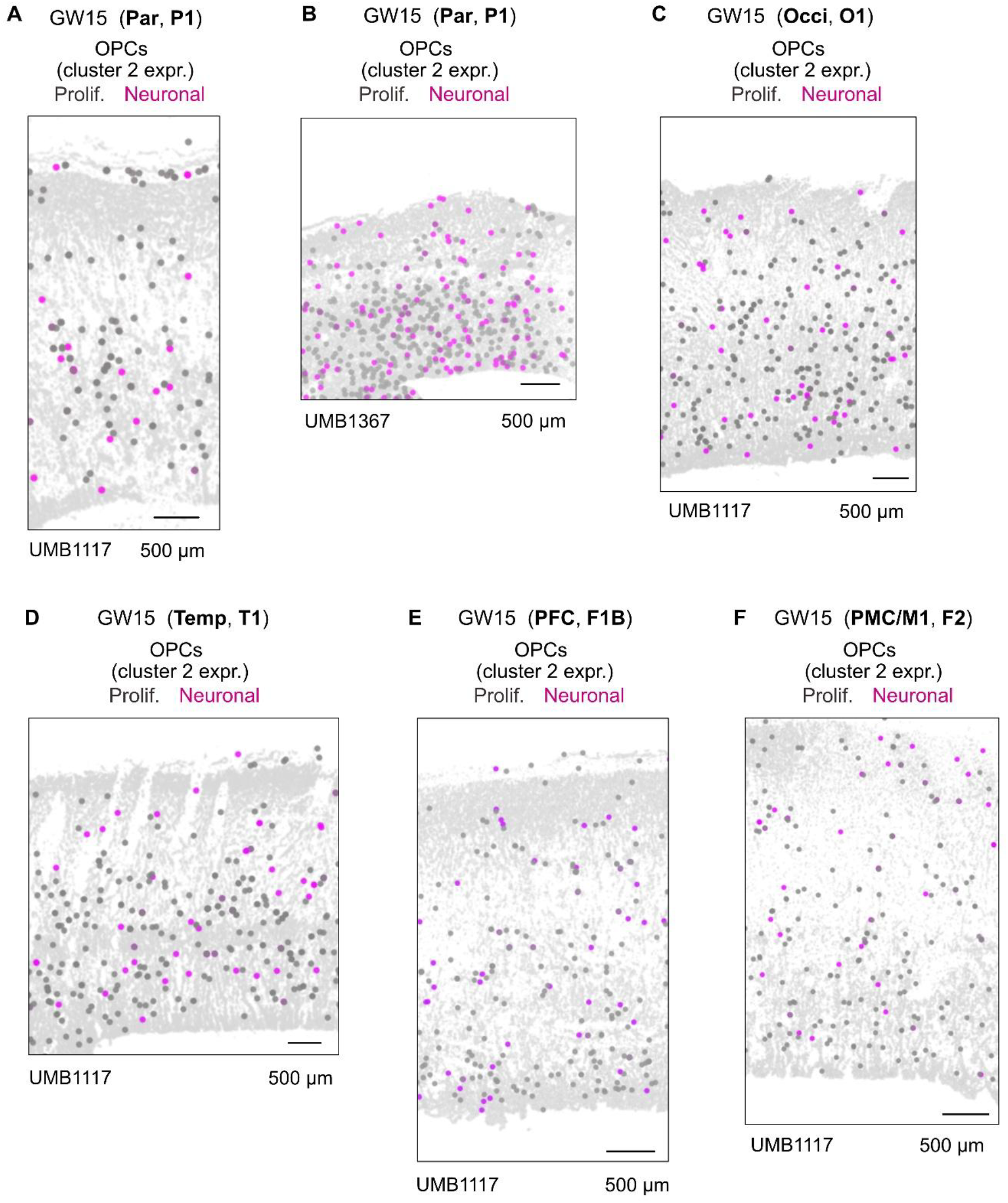
Laminar distribution of neuronal-OPC and proliferative-OPC cells in additional GW15 samples. **A to F**. Positions of neuronal-OPC (magenta) and proliferative-OPC (grey) cells in GW15 cortical sections, shown against all other cells in light grey. Sections span parietal cortex in two donors (**A**, UMB1117; **B**, UMB1367), occipital cortex (**C**), temporal cortex (**D**), prefrontal cortex (**E**) and premotor and primary motor cortex (**F**). Section identifiers are given in parentheses in each panel title. Cells were classified as in **Figure 6B**, taking the top 20% of OPCs by cluster 2 expression score as neuronal-OPC. Quantification of these conditions is given in **Figure 6D**. Scale bars are 500 µm.

**Supplementary Figure S10.**
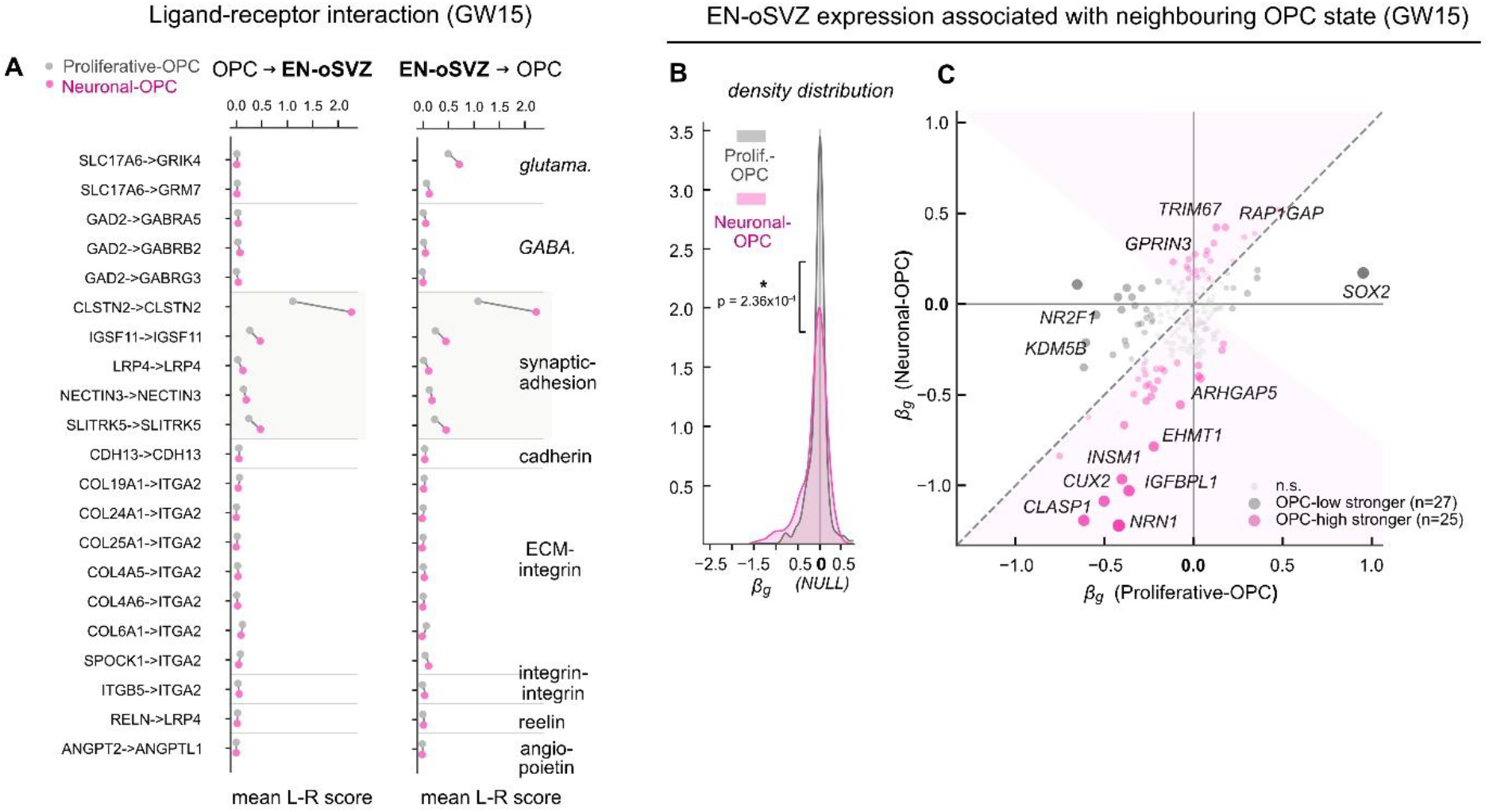
The association with neighbouring OPC state is weaker and less polarised at the EN-oSVZ interface. Analyses are performed as in Figure **6G-J**, with migrating excitatory neurons of the oSVZ (EN-oSVZ) rather than outer radial glia as the partner cell type. **A**. Ligand-receptor scores in neuronal-OPC and proliferative-OPC neighbourhoods, in both directions, grouped by interaction class. Among the synaptic-adhesion pairs, only *CLSTN2* scores significantly higher in neuronal-OPC neighbourhoods (*q* < 0.05, Mann-Whitney U), compared with four pairs at the oRG interface. No glutamatergic, GABAergic, cadherin, extracellular matrix or integrin pair differs between the two states. **B**. Distribution of the per-gene coefficient across the transcriptome in EN-oSVZ cells (n = 441,877), under each OPC state. The shift between states is smaller than at the oRG interface (Wasserstein distance 0.09 against 0.20; Kolmogorov-Smirnov *P* = 2.36 × 10^−4^). **C**. Coefficient under neuronal-OPC against coefficient under proliferative-OPC, per gene. Coloured points are significantly more strongly coupled to one state. Model specification, covariates and significance criteria are as in **Figure 6** and Methods. Per-gene coefficients are given in **Supplementary Table S8**.

